# Clinically-relevant altered antibiotic responses and mechanisms of β-lactam sensitization of MRSA in cystic fibrosis artificial sputum

**DOI:** 10.64898/2026.03.30.715424

**Authors:** Tyler J. Hamelin, Ali Molaeitabari, Marc R. MacKinnon, Tanya E. S. Dahms, Omar M. El-Halfawy

## Abstract

*Staphylococcus aureus* is the most common bacterial pathogen affecting pediatric patients with Cystic fibrosis (CF), a genetic disorder that causes thick mucus buildup in the lungs, providing a scaffold for chronic infections. Antibiotic treatment is typically guided by standard *in vitro* antimicrobial susceptibility testing (AST) in Mueller-Hinton broth (MHB), which does not represent the infection site in CF lungs. Notably, discordances between AST predictions and antibiotic therapeutic outcomes were reported in up to 50% of CF cases. To address this gap, we conducted ASTs against methicillin-resistant *S. aureus* (MRSA) in CF sputum-mimetic media compared with MHB, demonstrating ≥4-fold discordances across four of nine antibiotic classes. Most significantly, we observed unexpected β-lactam sensitization of MRSA strains (up to 128-fold) in CF sputum-like media, crossing the CLSI clinical breakpoint, suggesting this shift may alter therapeutic outcomes. Genome-wide screens and follow-up assays revealed underlying cell envelope remodelling and alterations to cell envelope stress responses. On the other hand, mucin binding to daptomycin may have led to an apparent 8-fold increase in resistance to this antibiotic in one of the CF sputum-like media. Overall, our AST results in CF sputum-mimetic conditions provide insights into bacterial responses during CF infections. Importantly, they suggest β-lactams may be effective in treating MRSA infections in CF patients, warranting further investigation in relevant *in vivo* systems.

## Introduction

Cystic fibrosis (CF) is a genetic disease caused by mutations in the gene encoding the CF transmembrane conductance regulator (CFTR), resulting in the accumulation of viscous mucus in the lungs (sputum) and on other mucosal surfaces^1^. Excess viscous mucus provides nutrients and a scaffold for microbes to reside and cause chronic respiratory tract infections, which are the primary cause of morbidity and mortality in CF patients^1^. Recently developed CFTR modulators, which correct the function defect of specific CFTR mutations, may greatly improve the prognosis of some CF patients^2^. However, CFTR modulators remain expensive, leaving many patients with limited access, do not correct all CFTR mutations, and do not improve the prognosis of patients with infections acquired before the initiation of the treatment^2–4^. As such, chronic infections continue to pose a serious threat to CF patients^4^.

*Staphylococcus aureus* is one of the most common causes of chronic cystic fibrosis respiratory tract infections and is the most common pathogen in children with CF, infecting 80% of CF patients aged 11 to 17^5, 6^. *S. aureus* is a Gram-positive pathogen that has developed multi-drug resistance, including to last resort antibiotics, and is on the World Health Organization’s list of antibiotic-resistant priority pathogens^7^. Persistent *S. aureus* airway infection is associated with several negative clinical outcomes, including deterioration of lung function, poorer nutritional parameters, and heightened airway inflammatory response in CF patients^8^. These negative outcomes have been particularly exacerbated by the emergence of methicillin-resistant *S. aureus* (MRSA) strains, which colonize more than 20% of CF patients and have contributed to lower rates of survival and compromised lung function^9, 10^.

Bacterial responses to antibiotics are often studied using antibiotic susceptibility testing (AST) assays standardized by globally recognized organizations such as the Clinical and Laboratory Standards Institute (CLSI)^11^. These assays use media such as Mueller-Hinton broth (MHB, composed of beef extract, casein hydrolysate, and starch^12^), which do not adequately represent the conditions at the CF lung infection site. The clinical outcome of antibiotic treatment in CF patients often does not correlate with antibiotic susceptibility results from these standard assays^13, 14^, and antibiotics recommended from these assays were successful in only half of the patients with respiratory tract infections^15^. Indeed, culture conditions and media composition dictate bacterial gene expression and dispensability profiles to meet bacterial survival requirements^16^. Culture media and conditions that mimic CF sputum have been developed^17–19^, such as artificial sputum medium (ASM)^17^ and synthetic cystic fibrosis medium-2 (SCFM-2)^19^. Notably, macromolecules such as mucin and DNA, which are more abundant in the matrix of respiratory tracts of CF patients during infection^17, 18^, may alter the activity of antimicrobials in the CF lung infection environment. Any such alterations are not detected under standard *in vitro* conditions but should be captured in infection-mimetic conditions. As expected, *S. aureus* showed altered antibiotic responses in media mimicking cystic fibrosis sputum^20^; however, the underlying mechanisms have not been investigated.

In this study, we used the CF sputum-like media, ASM and SCFM-2, to evaluate the antimicrobial susceptibility profile of MRSA compared with the standard MHB. We tested antibiotics from nine antibiotic classes; representatives from some of these classes had previously been tested in artificial CF sputum^20^, while others were not. The AST results against MRSA USA300, one of the most widely circulating community-acquired MRSA strains^21^, demonstrated ≥4-fold discordances in four of nine antibiotic classes tested between ASM and MHB. Most significantly, USA300 was unexpectedly more sensitive (up to 128-fold) to β-lactams in both CF sputum-like media, crossing the CLSI clinical breakpoint for oxacillin, suggesting this shift may alter therapeutic outcomes. Genome-wide assays and mechanistic investigations revealed that cell envelope remodelling and alterations to cell envelope stress responses underlie the unexpected hypersensitization to β-lactams. On the other hand, USA300 showed an apparent 8-fold increase in resistance to the lipopeptide daptomycin in ASM; however, potential interactions with mucin (e.g., direct binding of daptomycin to mucin) may contribute to this apparent shift. Together, this work addresses the shortcomings of MHB-based standard ASTs by uncovering altered antibiotic susceptibility patterns in CF sputum-like media compared with MHB, which may inform optimized antibiotic selection for CF patients. Mechanistic studies of the unexpected β-lactam sensitization of MRSA revealed potentially clinically relevant antibiotic-response determinants under CF conditions.

## Results and discussion

### MRSA susceptibility to antibiotics is altered in CF sputum-mimetic media

ASTs were performed using commonly used antibiotics from nine classes against USA300 in MHB and ASM (Table 1). Four of the nine antibiotics demonstrated significant alteration in minimum inhibitory concentration (MIC, ≥4-fold difference between ASM and MHB in all repeats). Colistin and daptomycin each displayed an 8-fold increase, and kanamycin displayed a 4-fold increase in MIC, while the β-lactam cefuroxime displayed a 128-fold reduction in MIC in ASM relative to MHB (Table 1). Notably, the observed 8-fold increase in colistin MIC is likely not clinically relevant as USA300 is already resistant to colistin, which is typically used to treat Gram-negative bacteria^22^. Additionally, the three antibiotics azithromycin, rifampicin, and tetracycline showed MIC shifts of 2–4-fold between ASM and MHB (Table 1). No or minimal (2-fold) MIC change between media was observed for ciprofloxacin and vancomycin (Table 1). These results match, for the most part, previously observed alterations in antibiotic susceptibility of another MRSA strain in ASM^20^. However, the differential susceptibility to colistin and daptomycin between MHB and ASM was not previously tested, whereas the tested MRSA strain (ATCC 33591) showed no change in the MIC of the carbapenem, meropenem, in both media^20^.

**Table 1.**
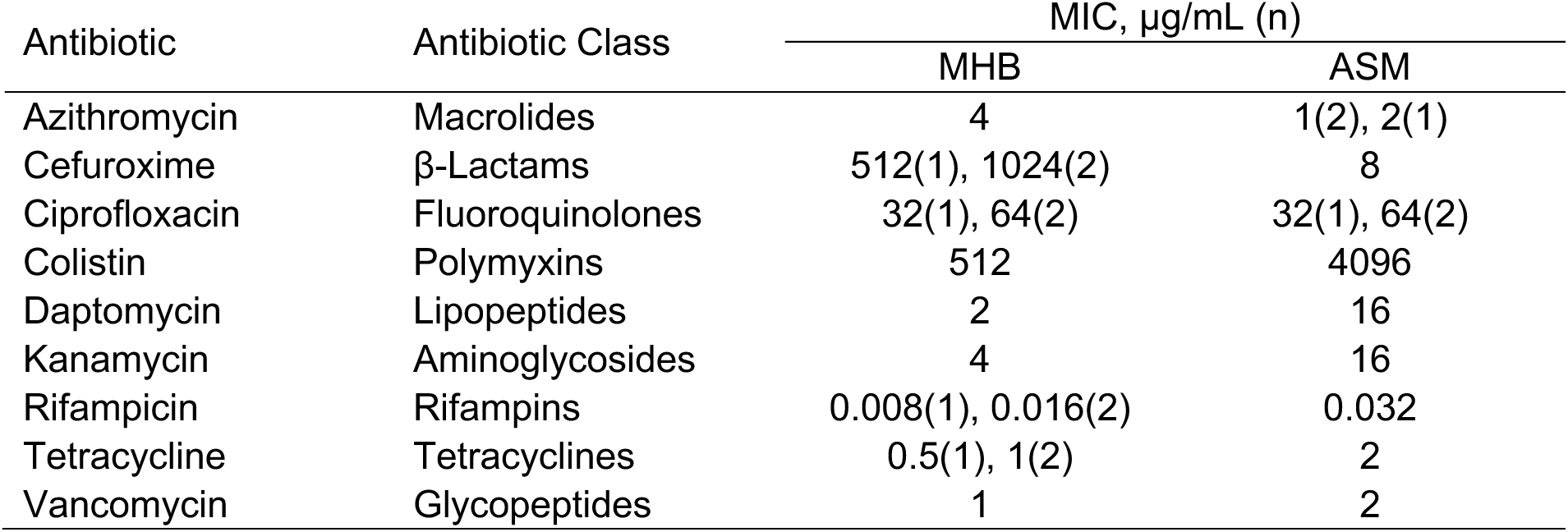
MIC of representative antibiotics against MRSA USA300 in ASM and MHB. MIC values are from three independent experiments (n = 3), each performed in triplicate. When MIC values across experiments did not match, the different values are reported with the corresponding n.

Together, the unexpected sensitization to cefuroxime and increased daptomycin resistance in MRSA USA300 were the most significant altered antibiotic-susceptibility phenotypes in ASM observed herein that may have potential clinical relevance; thus, we chose to further investigate these phenotypes. β-lactams, including cefuroxime, inhibit bacterial cell wall synthesis following the attachment of the β-lactam ring to penicillin-binding proteins, resulting in the interruption of peptidoglycan synthesis and subsequent cell death^23^. It is the most widely used antibiotic class with orally bioavailable options, but its use for *S. aureus* infections is prevented by the emergence of MRSA strains. Daptomycin is a cyclic lipopeptide with potent antimicrobial activity, typically used as a last resort against drug-resistant Gram-positive bacteria, such as MRSA^24^. Daptomycin exerts a calcium-dependent activity, inducing membrane depolarization and loss of intracellular components such as Mg^2+^, K^+^, and ATP^25^. Although daptomycin is typically not recommended for respiratory tract infections due to inhibition by pulmonary surfactants^26^, it has shown success in treating MRSA and other drug-resistant Gram-positive bacteria in pulmonary infections in *in vivo* models^27, 28^. Thus, it might be considered as a potential off-label therapeutic option for persistent MRSA infections in CF patients, or as an alternative for other systemic infections in these patients.

### Sensitization of MRSA strains to β-lactam antibiotics in CF sputum-mimetic media

Similar to the phenotype in ASM, we also observed a drastic sensitization of USA300 to cefuroxime in the other CF sputum-mimetic medium, SCFM-2, with an MIC similar to that in ASM (only 2-fold different; Table 2). We also tested oxacillin, another β-lactam used by CLSI to identify methicillin resistance for *S. aureus*. In ASM and SCFM-2, the MIC of oxacillin against USA300 was 32-fold lower than in MHB (Table 2), notably bringing the MIC below the CLSI breakpoint for resistance set at 4 μg/mL^29^. As such, ASTs performed in CF sputum-mimetic media suggest that β-lactam antibiotics may be effective for treating USA300 CF infections, contrary to standard AST recommendations that contraindicate these antibiotics for this MRSA strain. Similarly, testing a panel of isolates representing the most commonly circulating MRSA lineages in Canada (CMRSA isolates^30^) showed that all strains demonstrated a ≥4-fold decrease in β-lactam MIC in one or both of CF-mimetic media, except for CMRSA 1, 3, and 4 (Table 2). In addition, an MRSA isolate obtained from a pediatric CF patient in Regina, Saskatchewan, also showed increased β-lactam susceptibility in both ASM and SCFM-2, crossing the CLSI oxacillin breakpoint (Table 2). We also tested the susceptibility of two methicillin-sensitive *S. aureus* (MSSA) strains to β-lactams. Interestingly, we observed a ≥4-fold decrease in oxacillin MIC and only a 2-fold reduction in cefuroxime MIC against the well-characterized Newman strain in both CF sputum-like media relative to MHB (Table 2). However, we did not observe a significant shift in the susceptibility of an MSSA isolate obtained from a CF patient in Regina, Saskatchewan, to either β-lactam in the CF media (Table 2). This MSSA isolate was already susceptible to the tested β-lactams (e.g., its oxacillin MIC in MHB was 0.5 μg/mL, well below the CLSI breakpoint of 4 μg/mL^29^). These results suggest that no further sensitization to β-lactams may occur in CF artificial sputum media for extremely susceptible strains, although testing additional MSSA strains is required before reaching a definitive conclusion.

**Table 2.** MIC of cefuroxime and oxacillin against isolates representing the most commonly circulating MRSA and MSSA lineages in Canada and clinical isolates from CF patient. MIC values are from three independent experiments (n = 3), each performed in triplicate. When MIC values across experiments did not match, the different values are reported with the corresponding n.

| Strains | Lineage | Cefuroxime MIC, µg/mL (n) |  |  | Oxacillin MIC, µg/mL (n) |  |  |
| --- | --- | --- | --- | --- | --- | --- | --- |
|  |  | MHB | ASM | SCFM-2 | MHB | ASM | SCFM-2 |
| **MRSA** |  |  |  |  |  |  |  |
| JE2 | USA300 | 1024 | 8 | 16 | 32(2),<br>64(1) | 2 | 2 |
| CMRSA 1 | USA600 | >2048 | 2048 | >2048 | 1024 | 512 | 1024 |
| CMRSA 2 | USA100/800 | >2048 | 512(2),<br>1024(1) | 1024 | 512 | 128 | 256 |
| CMRSA 3 |  | >2048 | 1024 | 2048 | 512 | 256 | 512 |
| CMRSA 4 | USA200 | 2048 | 2048 | 2048 | 256 | 256 | 256 |
| CMRSA 5 | USA500 | 128 | 4 | 2 | 32 | 2 | 1(1), 2(2) |
| CMRSA 6 |  | >2048 | 2048 | 2048 | 1024 | 256 | 512 |
| CMRSA 7 | USA400/MW2 | 512 | 8 | 16(1),<br>32(2) | 64 | 16 | 16 |
| CMRSA 8 | EMRSA15 | 2048 | 32 | 128 | 128 | 64 | 64 |
| CMRSA 9 |  | 2048 | 512 | 1024 | 256 | 64 | 64 |
| CMRSA 10 | USA300 | 1024 | 8(2), 16(1) | 32 | 64 | 4 | 4 |
| CF clinical isolate* |  | 128(2),<br>256(1) | 8 | 8(2), 16(1) | 16(2),<br>32(1) | 2 | 2 |
| **MSSA** |  |  |  |  |  |  |  |
| Newman |  | 16 | 8 | 8 | 8 | 2 | 0.5 |
| CF clinical isolate** |  | 2 | 2 | 4 | 0.5 | 1 | 1 |

\**S. aureus* isolate from a CF patient, a 3-year-old boy from Regina, Saskatchewan, Canada
\*\**S. aureus* isolate from a CF patient from Regina, Saskatchewan, Canada

### Genomic screen reveals determinants of susceptibility to β-lactams in CF sputum-like media

Given the prevalence and magnitude of β-lactam sensitization across strains from multiple MRSA lineages, we next sought to determine the mechanism of cefuroxime susceptibility of MRSA in CF sputum-mimetic media using USA300 as a representative, being one of the most common community-associated MRSA strains. First, we performed genomic screens of the Nebraska transposon mutant library (NTML), a collection of sequence-defined transposon mutants that encompasses the majority of the nonessential genome of *S. aureus*^31^. We screened the NTML against different concentrations of cefuroxime in ASM and MHB (Fig. 1A) to address two possible hypotheses that the mechanism of the loss of cefuroxime resistance in ASM is mediated by: 1) factors involved in resistance that are expressed in MHB but not in ASM; and 2) factors that increase susceptibility to cefuroxime that are expressed only in ASM. To address the first hypothesis, we screened the NTML at concentrations corresponding to 1/16x and 1/8x of the wildtype MIC in MHB, indentifying 57 mutants with reduced growth relative to the wildtype and the interquartile mean (other unrelated mutants), hence may harbour mutations in genes potentially involved in increased β-lactam resistance in MHB (Fig. 1B). To address the second hypothesis, we screened the NTML at cefuroxime concentrations corresponding to 4x and 8x MIC in ASM, identifying 111 mutants with increased growth relative to the wildtype and the interquartile mean (Fig. 1C). Follow-up dose-response and MIC assays against the identified 57 and 111 mutants revealed that 14 mutants exhibited a ≥4-fold sensitization to cefuroxime relative to the wildtype in MHB with no or minimal reduction in ASM (Table 3, Supplementary Fig. 1) while 10 mutants showed a ≥4-fold increase in resistance to cefuroxime relative to the wildtype in ASM with no or minimal increase in MHB (Table 3, Supplementary Fig. 2).

**Figure 1.**
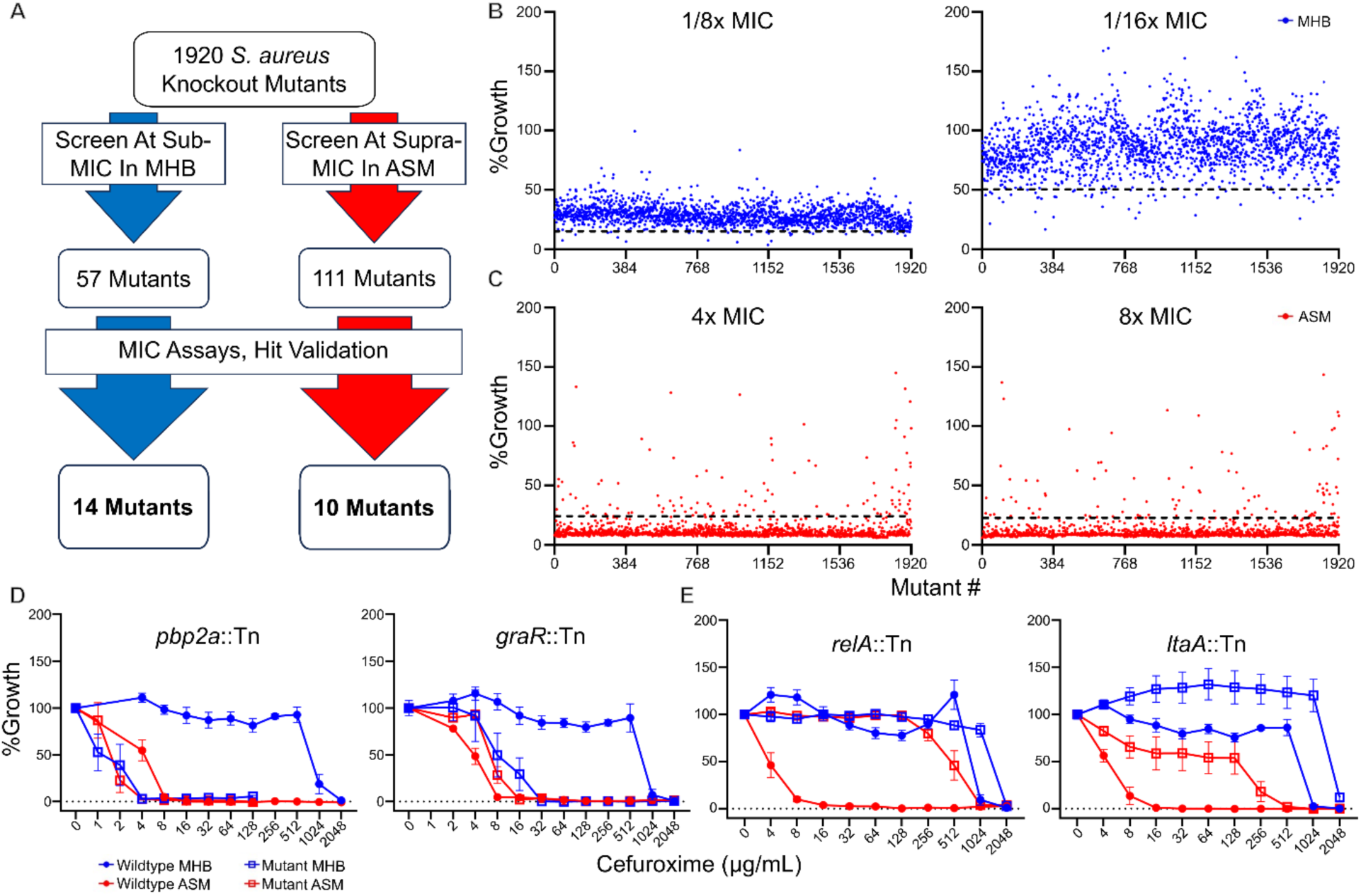
A genomic screen identifies putative determinants of USA300 sensitivity to cefuroxime in ASM. A) A flow chart outlining the steps taken to filter and identify mutants of putative determinants. B) Index plots for the NTML screened at sub-MIC concentrations of cefuroxime in MHB. C) Index plots for the NTML screened at supra-MIC concentrations of cefuroxime in ASM. The dotted lines represent the cutoff below which (B) and above which (C) mutants were considered potential putative determinants selected for further follow-up. D) Dose-response growth curve for *pbp2a*::Tn and *graR*::Tn, two mutants identified from screening in MHB. E) Dose-response growth curve for *relA*::Tn and *ltaA*::Tn, two mutants identified from screening in ASM. Growth curves represent n=6 from three independent experiments.

**Table 3.**
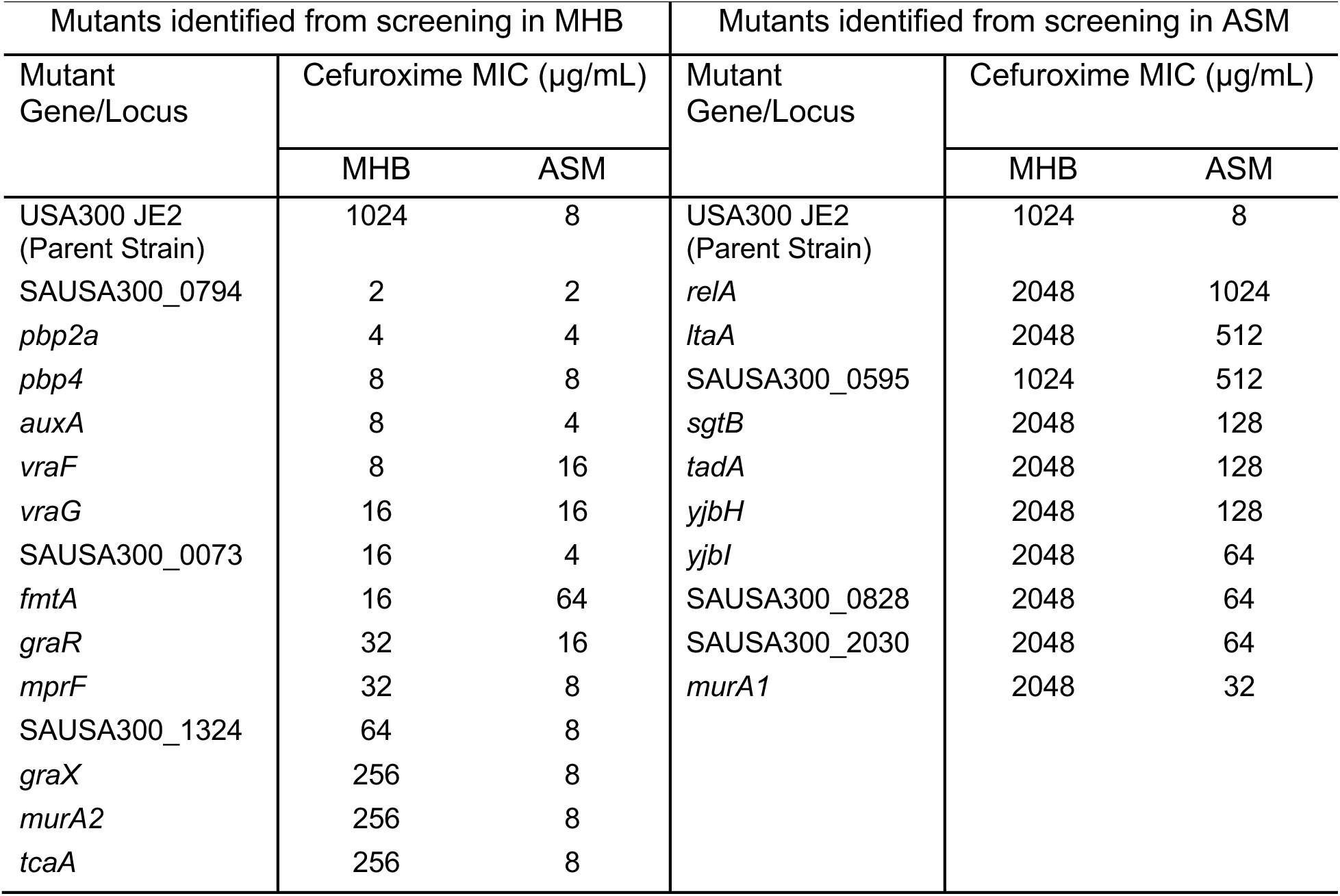
MIC of cefuroxime against mutants identified from screening the NTML in MHB and ASM. MICs represent the average of three independent experiments (n = 3), each performed in duplicate. Corresponding growth curves for MICs are shown in Supplementary figures 1 and 2.

Most of the identified mutants harbour mutations that disrupt processes or factors broadly related to the cell envelope and cellular stress responses. Mutants related to the cell envelope include those related to peptidoglycan (PG; *pbp2a*, *pbp4*, *murA1, murA2, lytH*, *sgtB*, and *yjbH*), wall teichoic acid (WTA; *fmtA* and *tcaA*), and lipoteichoic acid (LTA; *ltaA* and *auxA*) and those localized to the cell envelope (SAUSA300_0073, SAUSA300_1324, SAUSA300_2030). The mutants related to cellular stress responses are those within the cell envelope stress response GraXRS/VraFG five-component system (*graR, graX, vraF*, *vraG,* and *mprF*) and the stringent response (*relA*). The remaining four mutants of the putative determinants are two that have been previously found to influence β-lactam susceptibility [SAUSA300_0595^32^ and SAUSA300_0794^33^], and two not previously linked to β-lactam resistance [*tadA,* encoding a tRNA adenosine deaminase^34^, and SAUSA300_0828, encoding a 5’-nucleotidase family protein^35^]. Importantly, many of the identified genetic determinants related to the different cell envelope structures (PG, WTA, and LTA) and stress response (e.g., GraXRS/VraFG) are genetically interconnected as shown in genetic lethal interaction analyses^36^. As such, we next set out to characterize alterations in the cell envelope and stress response and assess their implications for the unexpected β-lactam hypersensitivity of MRSA USA300 in CF-mimetic media.

### Alterations to peptidoglycan are apparent under CF sputum-mimetic conditions

First, we investigated potential alterations in penicillin-binding proteins (PBPs) in CF-mimetic media, given that PBPs are the targets of β-lactams and that the primary resistance mechanism of MRSA USA300 is PBP2a, encoded by *pbp2a*/*mecA*^37^. Based on the observed phenotypes of *pbp2a*::Tn and *pbp4*::Tn (Table 3, Supplementary Fig. 1), we hypothesized that USA300 has an overall decreased abundance of PBPs when cultured in the CF sputum-mimetic media. To assess the abundance of PBPs, we isolated membrane fractions from cultures grown in MHB and SCFM-2. We used SCFM-2 in this and other assays requiring cell isolation at mid-log phase (when cell density is relatively low) as SCFM-2 contains 1/4x the amount of mucin in ASM, enabling the collection of the required cell density from the SCFM-2 cultures without excessive cell loss that occurs while eliminating the higher mucin content in ASM. We used a fluorescent penicillin (BOCILLIN^TM^ FL) to visualize all PBPs resolved on SDS-PAGE, except for PBP2a, which has low binding affinity to this penicillin (Fig. 2A, Supplementary Fig. 3). Band quantification revealed no statistically significant alteration in the abundance of PBP1, but a ∼27%, ∼56%, and ∼42% reduction in PBP2, PBP3, and PBP4, respectively, in SCFM-2 compared to MHB (Fig. 2B-E). Notably, PBP2 is an essential gene and is thus not represented in the NTML. Similarly, Western blots using anti-PBP2a antibody (Fig. 2F, Supplementary Fig. 4) showed a 20% reduction in PBP2a in SCFM-2 compared to MHB (Figure 2G). Together, the decreased abundance of four PBPs (2, 2a, 3, and 4) is likely a major contributing factor for the hypersensitivity of USA300 to β-lactams in CF sputum-like conditions.

**Figure 2.**
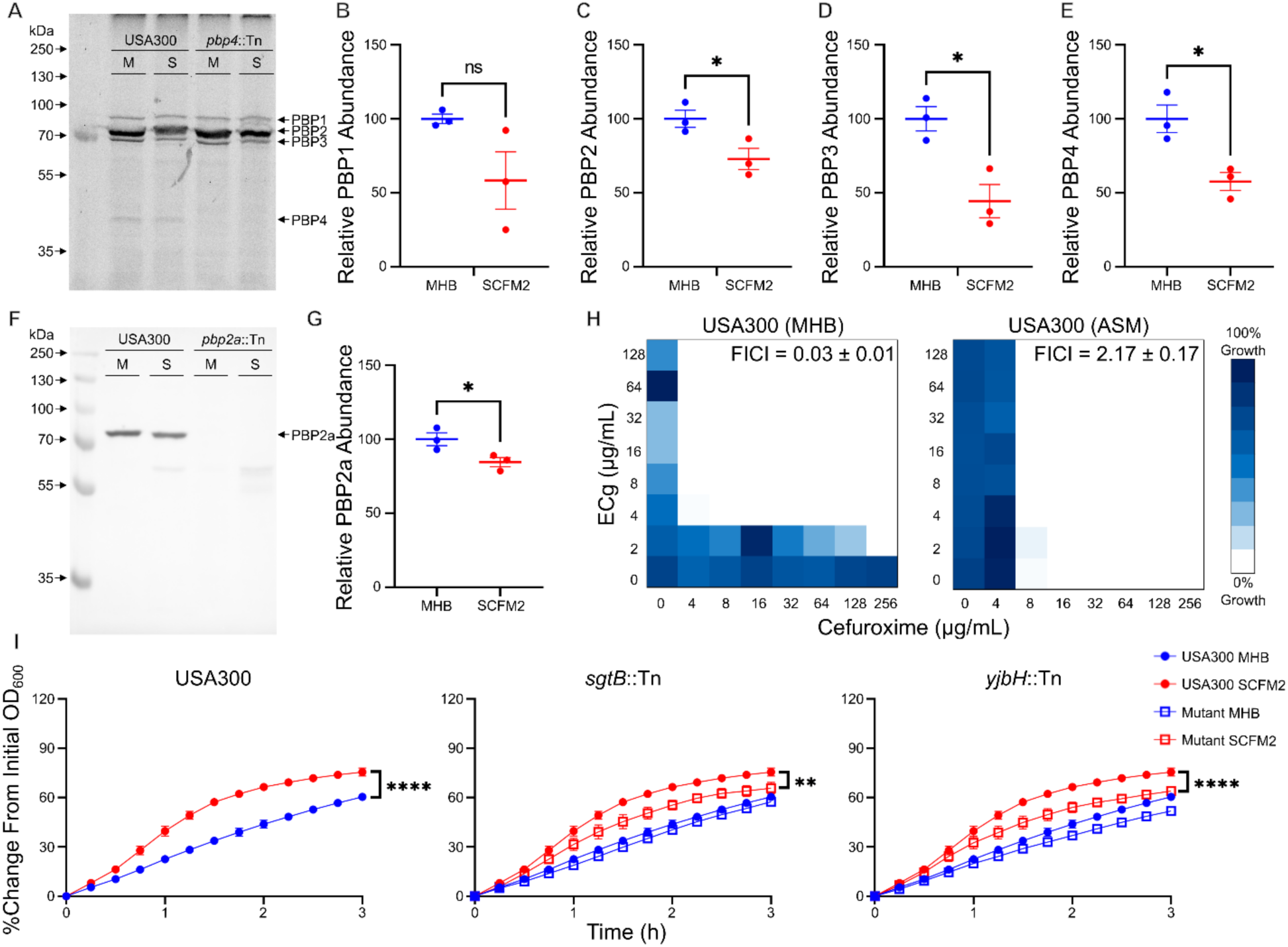
Decreased abundance of PBPs and alterations in PBP activity in CF sputum-mimetic media contribute to cefuroxime sensitivity. A) Representative BOCILLIN FL-labelled PBP gel from membrane fractions isolated from USA300 and *pbp4*::Tn grown in MHB (M) or SCFM-2 (S). Gels for all isolations (n=3) are shown in Supplementary Fig. 3. Quantification of B) PBP1, C) PBP2, D) PBP3, and E) PBP4 from USA300 cultured in SCFM-2 (red) relative to MHB (blue), n=3. F) Representative Western blot for PBP2a using anti-PBP2a antibody on membrane fractions isolated from USA300 and *pbp2a*::Tn cultured in MHB (M) or SCFM-2 (S). Blots for all isolations (n=3) are shown in Supplementary Fig. 4. G) Quantification of PBP2a from USA300 cultured in SCFM-2 (red) relative to MHB (blue), n=3. H) Representative checkerboard assays of *S. aureus* USA300 for cefuroxime in combination with epicatechin gallate (ECg) performed in MHB or ASM, FICI shown as mean ± SEM (n=3). I) Lysostaphin-induced lysis turbidimetric assays performed against *S. aureus* USA300, *sgtB*::Tn, and *yjbH*::Tn cultured in either MHB or SCFM-2 (n = 9 from three independent experiments, shown as mean ± SEM). Significant differences were identified using Welch’s t-test (B, C, D, E, and G) or two-way ANOVA and the Tukey post hoc test (I). p<0.0001(**\*\*\*\***), p<0.001(**\*\*\***), p<0.01(**\*\***), and p<0.05(**\***).

Next, we investigated whether disruption of PBP coordination would confer the cefuroxime susceptibility observed in ASM. Epicatechin gallate (ECg) is a polyphenol found in green tea that reverses β-lactam resistance by delocalizing PBP2 from the division septum^38^. To this end, we tested ECg in combination with cefuroxime against USA300 in both MHB and ASM (Fig. 2H). As expected, ECg synergized with cefuroxime in MHB against USA300 (fractional inhibitory concentration index [FICI] = 0.03 ± 0.01). However, this synergy was lost in ASM (FICI = 2.17 ± 0.17), suggesting that PBP coordination and PBP2 localization were already disrupted in the CF medium.

The activity of PBPs is critical for maintaining PG crosslinking. Therefore, we assessed the rates of lysostaphin-induced lysis of cells harvested from MHB or SCFM-2 cultures. As expected, lysis rates of wild-type cells from MHB cultures were slower than those from SCFM-2 (Fig. 2I). *sgtB*::Tn and *yjbH*::Tn, which have previously been reported to have increased PG crosslinking^39, 40^ and were more resistant to cefuroxime only in ASM (Table 3, Supplementary Fig. 2), showed slower lysis rates than the wildtype in SCFM-2. Together, our results suggest that there are overall reductions in PBP abundance, PBP activity, and PG crosslinking in CF sputum-like media, which may contribute to cefuroxime sensitivity.

### Alterations in teichoic acids in CF sputum-mimetic conditions

Next, we evaluated alterations to the cell envelope appendages WTA and LTA, which play roles in coordinating PG synthesis and β-lactam resistance^41^. WTA and LTA content and modifications contribute, along with other cell envelope modifications, to cell surface hydrophobicity^42–44^, where mutants deficient in WTA or with a drastic reduction in LTA exhibit significantly increased hydrophobicity^42, 43^. Hence, we measured the cell surface hydrophobicity of wildtype USA300, using the microbial adherence to hydrocarbon (MATH) method^45^. The wildtype USA300 cells cultured in ASM were significantly more hydrophobic than those cultured in MHB (Fig. 3A), suggesting potential reduced TA content in ASM.

**Figure 3.**
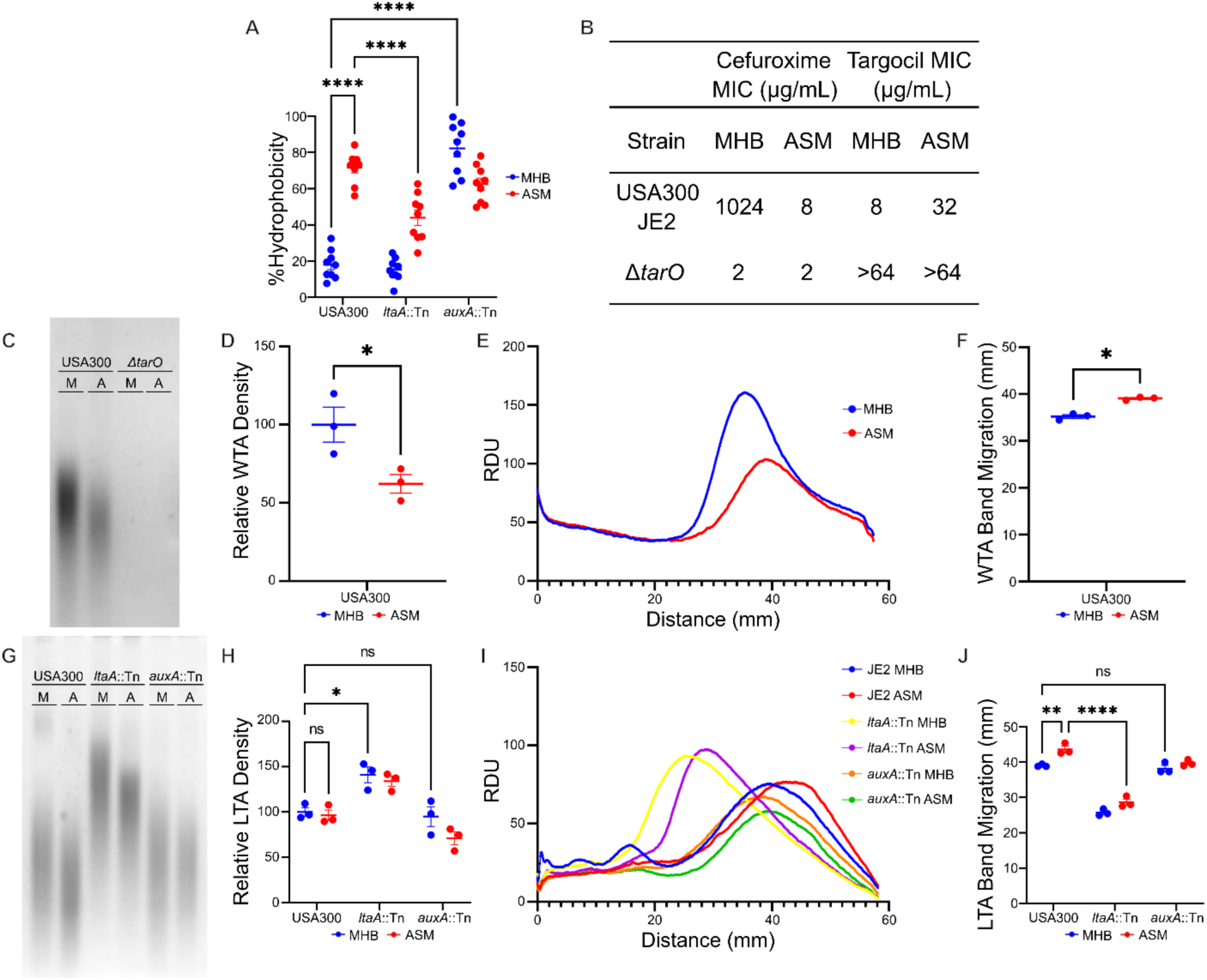
Teichoic acids are modified in CF sputum-mimetic media. A) Cell surface hydrophobicity assay performed on USA300, *ltaA*::Tn, and *auxA*::Tn cultured in either MHB (blue) or ASM (red), n=9 from three independent experiments. B) MICs for cefuroxime and targocil against USA300 JE2 and Δ*tarO* performed in MHB or ASM, from three independent experiments. C) Representative PAGE gel for crude WTA isolated from USA300 or Δ*tarO* cultured in MHB (M) or ASM (A). Gels for all isolations (n=3) are shown in Supplementary Fig. 5. D) Quantification of the overall band intensity of crude WTA isolated from USA300 cultured in ASM (red) relative to MHB (blue), shown as mean ± SEM. E) Relative band migration of crude WTA isolated from USA300 cultured in MHB or ASM, represented as the mean relative pixel density units (RDU), n=3. F) Analysis of band migration for crude WTA isolated from USA300 cultured in MHB (blue) and ASM (red), shown as mean of the distance corresponding to the peaks shown in E ± SEM. G) Representative PAGE gel for crude LTA isolated from USA300, *ltaA*::Tn, or *auxA*::Tn cultured in either MHB (M) or ASM (A). Gels for all isolations (n=3) are shown in Supplementary Fig. 6. H) Quantification of crude LTA isolated from USA300, *ltaA*::Tn, or *auxA*::Tn cultured in either MHB (blue) or ASM (red), shown as mean ± SEM. I) Relative band migration of crude LTA isolated from USA300, *ltaA*::Tn, or *auxA*::Tn cultured in either MHB or ASM, represented as the mean pixel density ± SEM, n=3. J) Analysis of band migration of crude LTA isolated from USA300, *ltaA*::Tn, or *auxA*::Tn cultured in either MHB (blue) or ASM (red), shown as the mean distance corresponding to the peaks shown in I ± SEM. Significant differences were identified using Welch’s t-test (D and F) or two-way ANOVA and the Tukey post hoc test (A, H, and J). p<0.0001(**\*\*\*\***), p<0.001(**\*\*\***), p<0.01(**\*\***), and p<0.05(**\***).

To evaluate alterations in WTA, we tested a knockout of *tarO*, which encodes an *N*-acetylglucosamine-1-phosphate transferase, the first committed step in WTA production^41^. As a *tarO*::Tn is not present in the NTML, it was not represented in our screen results. A CMRSA10 Δ*tarO* (herein referred to as Δ*tarO*) was 512-fold more susceptible to cefuroxime in MHB relative to the wildtype and not further sensitized to the β-lactam in ASM (Fig. 3B), mirroring the primary determinants identified above. Next, we used targocil, an inhibitor of the WTA flippase TarG^41^, to determine whether culturing in ASM or SCFM-2 disrupts WTA biosynthesis. Targocil is bactericidal; however, disrupting genes encoding the early steps of WTA synthesis (*tarO* and *tarA*) or inhibiting their products (blocking WTA biosynthesis) antagonizes targocil. In ASM, USA300 was 4-fold more resistant to targocil in ASM relative to MHB (MIC values of 32 and 8 μg/mL, respectively; Fig. 3B). In comparison, Δ*tarO* displayed a targocil MIC >64 μg/mL in both MHB and ASM (Fig. 3B), suggesting that WTA biosynthesis is, at least partially, disrupted in the CF sputum-like media. In addition, we isolated WTA polymers from cells grown in MHB and ASM, separated them using PAGE, and visualized their bands using alcian blue staining (Fig. 3C, Supplementary Fig. 5). We quantified the WTA PAGE band intensities, revealing a significant reduction in WTA density from cells grown in ASM compared to MHB (Fig. 3D). Further, WTA from cells grown in ASM demonstrated a slightly faster migration of WTA on PAGE (Fig. 3E,F), suggesting a shorter polymer length. Δ*tarO* was included as a negative control due to its inability to produce WTA, revealing no WTA bands in either medium, as expected (Fig. 3C). Together, this data shows alterations in WTA when MRSA USA300 is cultured in the CF sputum-like media.

Similarly, we isolated, separated, and visualized LTA from cells grown in MHB and ASM to evaluate alterations to LTA in ASM (Fig 3G, Supplementary Fig. 6). While there was no significant alteration in the overall band intensity of LTA in wildtype USA300 between MHB and ASM (Fig. 3G,H), LTA from cells grown in ASM demonstrated a statistically significant slightly faster migration of LTA, possibly suggesting a shorter polymer length (Fig. 3I,J). Controlling LTA polymer length is critical for cell envelope integrity and β-lactam response^46^; thus, shorter LTA polymers may contribute to β-lactam hypersensitivity in ASM. Notably, *ltaS,* which encodes LTA synthase, is an essential gene^47^; hence, we could not include a mutant with complete loss of LTA content. Instead, we tested two LTA-related mutants, *ltaA*::Tn and *auxA*::Tn^46, 48^. Relative to the wildtype strain, *ltaA*::Tn is more resistant in ASM (Table 3, Supplementary Fig. 2), and *auxA*::Tn is more susceptible in MHB (Table 3, Supplementary Fig. 1). *ltaA*::Tn and *auxA*::Tn showed increased and similar quantities of LTA compared to the wildtype, respectively (Fig. 3H), consistent with previous literature^46, 48^. Of note, an increased release of LTA in the

supernatant was previously shown in *auxA*::Tn^48^. Further, LTA from *ltaA*::Tn and *auxA*::Tn demonstrated slower and similar migration relative to the wildtype on PAGE gel, respectively (Fig. 3I,J). Notably, *ltaA*::Tn showed reduced hydrophobicity in ASM, whereas *auxA*::Tn exhibited increased hydrophobicity in MHB compared to the parent strain (Fig. 3A). Together, these results suggest that TA alterations, including reduced WTA content and LTA polymer length, are associated with increased β-lactam susceptibility in CF sputum-mimetic media.

### The GraXRS/VraFG stress response is not induced in CF sputum-mimetic conditions

GraXRS/VraFG is a five-component system that responds to cell envelope stressors such as cationic antimicrobial peptides^49^, with GraR, VraF, VraG, and MprF shown to be involved in β-lactam resistance^50^. *mprF,* whose product influences cell surface charge through L-lysine modification of phosphatidylglycerol, is positively regulated by GraXRS, and colistin, a cationic antimicrobial peptide, strongly induces the system^49, 50^. Thus, we monitored *mprF* expression through a transcriptional fusion with *luxABCDE* in a luciferase expression assay in tryptic soy broth (TSB), a standard medium, compared to ASM. In TSB, the expression of *mprF* was induced in the presence of colistin as expected, indicating a functional GraR (Fig. 4A). In ASM, colistin failed to induce *mprF* expression above the untreated control, a result similar to those observed in a Δ*graR* mutant (Fig. 4B). We tested the promoter of *ilvD*, encoding a protein involved with the synthesis of branched chain amino acids, as a negative control^50^, showing no changes in luciferase signal above baseline (Fig. 4C). Together, these results indicate that GraXRS is not induced by cell envelope stress under CF sputum-mimetic conditions.

**Figure 4.**
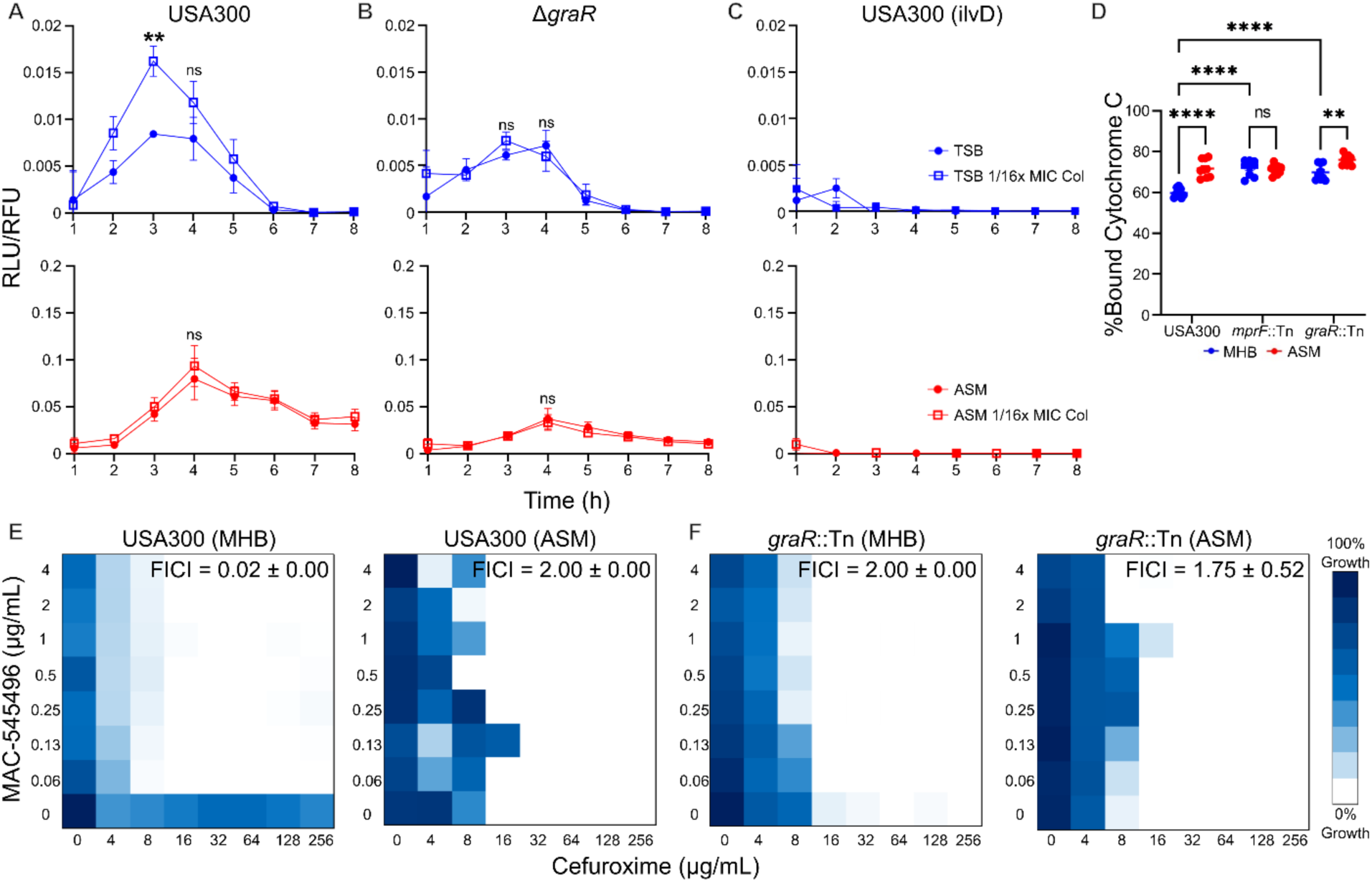
Disruption of GraXRS/VraFG in CF sputum-mimetic conditions contributes to β-lactam sensitivity. Monitoring luciferase production under the control of the promoter for *mprF* in A) USA300 and B) ΔgraR, and C) under the control of the promoter for *ilvD* in USA300 as a control, n=9 from three independent experiments. Higher RLU signals were observed in ASM due to the presence of egg yolk emulsion (see the Supplementary Methods and the associated data for details). D) Cytochrome C binding assays performed on USA300, *mprF*::Tn, or *graR*::Tn cultured in either MHB (blue) or ASM (red), n=9 from three independent experiments. Representative checkerboards for cefuroxime performed in combination with MAC-545496 against E) USA300 and F) *graR*::Tn in MHB or ASM, FICI values reported as mean ± SEM, n=3. Significant differences were identified using two-way ANOVA and the Tukey post hoc test. p<0.0001(**\*\*\*\***), p<0.001(**\*\*\***), p<0.01(**\*\***), and p<0.05(**\***).

Further, we tested the binding of cytochrome C, a positively charged protein, to cells cultured in MHB compared to ASM. Disruptions in *graR and mprF* can lead to an increase in the overall net negative charge of *S. aureus*, leading to increased binding of cells to cytochrome C^51–53^. Indeed, cells grown in ASM were able to bind cytochrome C significantly more than cells grown in MHB (Fig. 4D). Further, *mprF*::Tn and *graR*::Tn also had significantly greater binding to cytochrome C, similar to that of USA300 grown in ASM. Of note, a GraXRS/VraFG disruption-related increase in the overall net negative charge may be mediated by the lack of induction of the *dlt* operon, whose products catalyze the D-alanylation of teichoic acids to mask their negative charge^44^, regulated by this five-component system^49^. However, such *dlt*-mediated effect might be offset by the overall reduction in WTA content when the wild-type is cultured in the CF sputum-like medium (Fig. 3C,D). Additionally, we tested MAC545496, an inhibitor of GraR that has been shown to synergize with β-lactams^50^, in combination with cefuroxime in both MHB and ASM. As expected, a GraR-dependent synergy between MAC545496 and cefuroxime was observed in MHB (Fig. 4E, FICI = 0.02 ± 0.00). This synergistic interaction was lost in ASM (FICI = 2.00 ± 0.00), mimicking the phenotype of *graR*::Tn (Fig. 4F, FICI = 2.00 ± 0.00 and 1.75 ± 0.52 in MHB and ASM, respectively). Together, the lack of *mprF* promoter induction, alteration in cell surface charge, and loss of MAC545496-cefuroxime synergy illustrate the loss of stress-mediated induction of GraR and its regulon, and thus the disruption of the GraXRS/VraFG system, in CF sputum-mimetic conditions.

### Induction of stringent response leads to reversal of β-lactam sensitivity in the CF sputum-like medium

Stringent response is controlled by the production of the alarmones (p)ppGpp in response to cellular stress, including β-lactam treatment, which leads to increased PBP2a expression^54^. Mutants lacking activity of both RelP and RelQ, two (p)ppGpp synthases expressed in conditions of cell wall stress, are more sensitive to cell wall-active antibiotics^54^. In contrast, mutations in RelA that result in constant activation of the stringent response, such as a truncation of the C-terminal domain^55^, led to increased β-lactam resistance^56^, where RelA includes (p)ppGpp synthesis and hydrolysis domains and a C-terminal domain that regulates the dominant activity^55^. Since the *relA*::Tn mutant, which has an insertion in the C-terminal regulatory domain^31^, reversed the cefuroxime sensitization of USA300 in CF sputum-like medium (Table 3, Supplementary Fig. 2), we sought to assess the potential contribution of the stringent response in this phenotype. To that end, we measured the intracellular level of (p)ppGpp in USA300 wildtype and *relA*::Tn in both MHB and SCFM-2 using the fluorescent probe PyDPA^57^ with and without exposure to cefuroxime or mupirocin, a stringent response activator^58^. As expected, *relA*::Tn showed a significant increase in (p)ppGpp in MHB and SCFM-2 relative to the wildtype (p<0.001 and p<0.0001, respectively; Fig. 5A), which matches with the increased cefuroxime resistance of that mutant (Table 3, Supplementary Fig. 2). Interestingly, while cefuroxime treatment only slightly, albeit statistically insignificantly, increased (p)ppGpp in MHB, it significantly decreased the level of the alarmones in SCFM-2 (Fig. 5A), which may indicate a lack of stringent response induction that matches with the β-lactam sensitization. Mupirocin treatment was included as a positive control and resulted in increased (p)ppGpp in both MHB and SCFM-2 relative to the wildtype (Fig. 5A).

**Figure 5.**
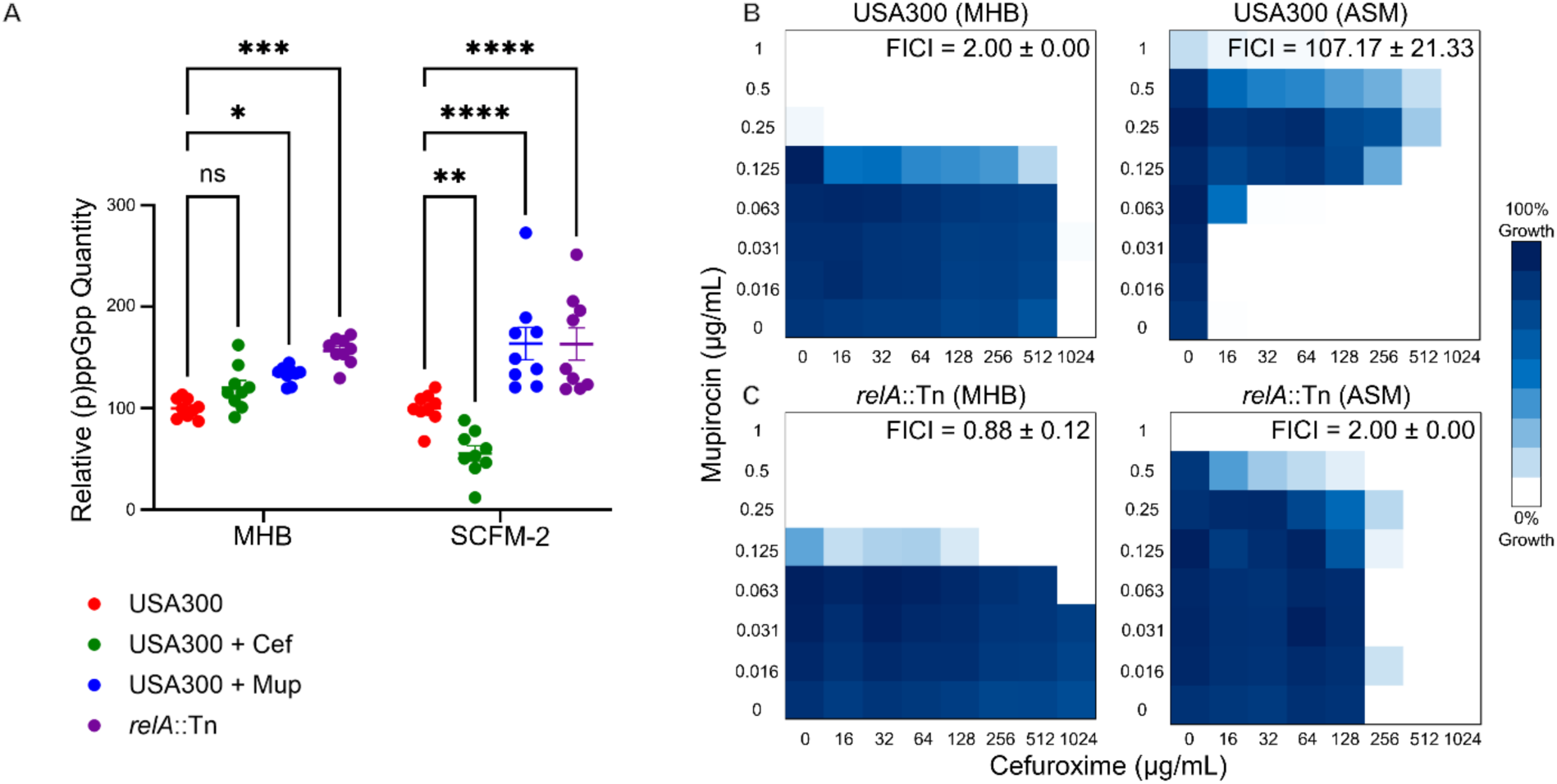
Stringent response in response to β-lactam treatment is altered in CF sputum-mimetic media. A) Quantification of (p)ppGpp produced by USA300 (with or without treatment with cefuroxime or mupirocin) and *relA*::Tn cultured in MHB or SCFM-2; mean ± SEM, n=9 from three independent experiments. B,C) Representative cefuroxime-mupirocin checkerboard assays in MHB or ASM against A) USA300 or B) *relA*::Tn; FICI values reported as mean ± SEM (n=3). Significant differences were identified using two-way ANOVA and the Tukey post hoc test. p<0.0001(**\*\*\*\***), p<0.001(**\*\*\***), p<0.01(**\*\***), and p<0.05(**\***).

To assess whether changes in the induction of stringent response underlie the β-lactam sensitization in the CF sputum-like media, we tested a combination of mupirocin and cefuroxime in both MHB and ASM. There was no significant interaction between the two antibiotics against USA300 in MHB (FICI = 2.00 ± 0.00), while in ASM there was a significant antagonistic interaction against the wild-type (FICI = 107.17 ± 21.33), leading to a reversal of the hypersensitivity to β-lactams (Fig. 5B). In comparison, no antagonism was observed against *relA*::Tn, which already has stringent response induced, in both MHB and ASM (FICI = 0.88 ± 0.12 and 2.00 ± 0.00; Fig. 5C). Together, these results suggest the lack of stringent response induction or β-lactam-mediated suppression thereof under CF-like conditions play a role in β-lactam sensitization in the CF sputum-like media, possibly through regulating PBP2a expression, whose abundance is reduced in CF media.

### Atomic force microscopy further supports alterations at the cell envelope under CF sputum-like conditions

We further assessed cells grown in CF sputum-mimetic media and their envelopes using atomic force microscopy (AFM; Fig. 6A) to compare surface topography, roughness, cell size, elasticity, and adhesion (Fig. 6B-C, Supplementary Fig. 7). We observed a significant increase in surface roughness and cell diameter in cells grown in SCFM-2 compared to those grown in MHB (Fig. 6D, E), but no significant alteration in cell height (Fig. 6F). The surface of USA300 cells grown in SCFM-2 was more compliant relative to MHB, requiring less force to deform (Fig. 6G). Adhesion between the negatively charged silicon nitride AFM tip and the cell surface was reduced for cells grown in SCFM-2 compared to MHB (Fig. 6H), consistent with differences in cell surface charge observed in the cytochrome C binding assay (Fig. 4D). Together, this data provides further evidence for cell envelope-related alterations under CF-mimetic conditions, which underpins the hypersensitivity to β-lactams under such conditions.

**Figure 6.**
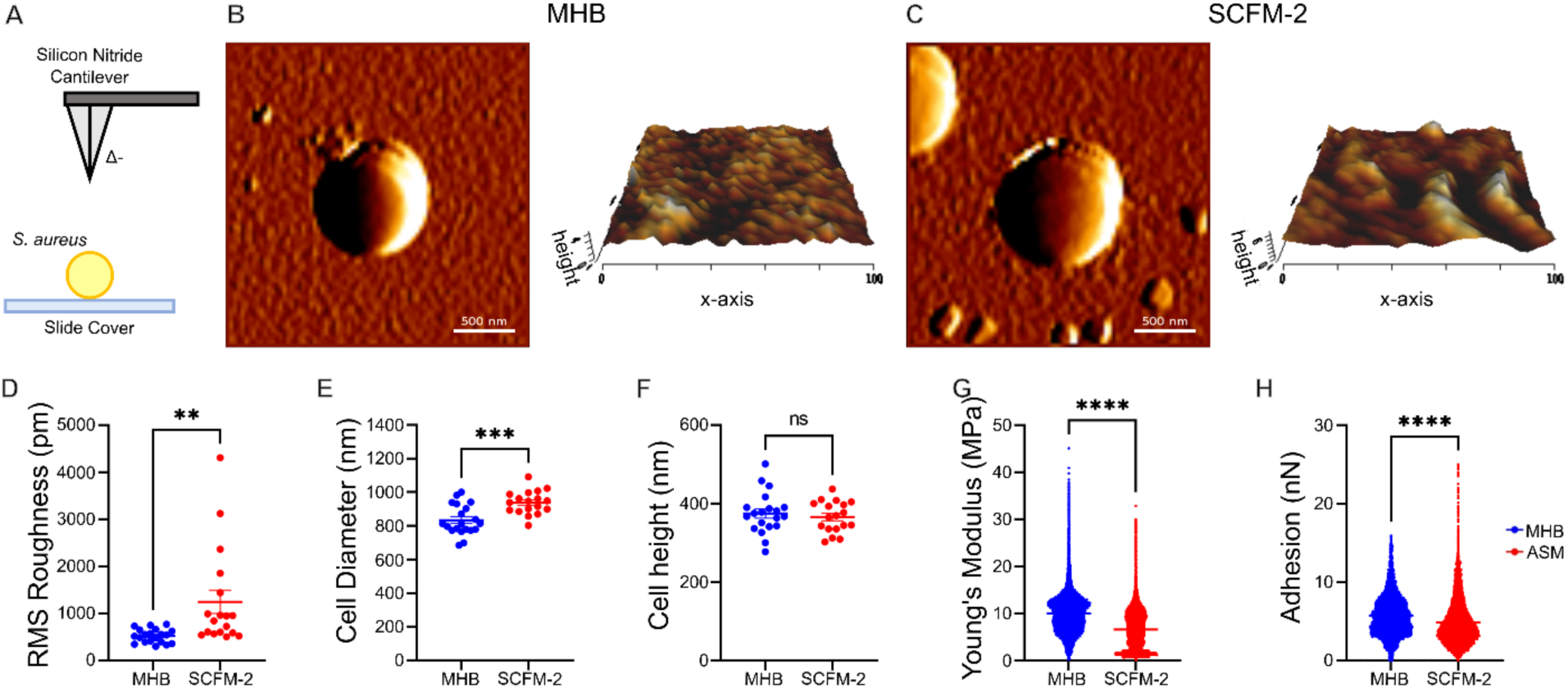
Atomic force microscopy of USA300 in MHB versus CF sputum-mimetic media. A) A schematic diagram of the AFM probe used for imaging. B, C) Representative 2D and 3D images of cell surface topography for cells cultured in B) MHB and C) SCFM-2. All images used for quantitative analysis are shown in Supplementary Fig. 7. D) Quantification of surface roughness as root mean square (RMS) deviation from mean height for the flattest 100 nm x 100 nm region in the center of the cell, within a 400 nm x 400 nm scanning area. E, F) Cell diameter and cell height of USA300 grown in MHB (blue, n=20) or SCFM-2 (red, n=18). G-H) Viscoelasticity of and adhesion to the AFM tip for USA300 cultured in MHB (n=295752) and SCFM-2 (n=218883). Images represent a minimum of 18 cells from three biological replicates. Significant differences were identified using Welch’s t-test, with significance denoted by asterisks: p<0.0001(**\*\*\*\***), p<0.001(**\*\*\***), p<0.01(**\*\***), and p<0.05(**\***).

### Increased β-lactam susceptibility is influenced by multiple CF sputum media components

We sought to investigate which ASM component may be responsible for altering *S. aureus* β-lactam susceptibility. First, we titrated individual ASM components into MHB while determining the cefuroxime MIC in checkerboard format. Mucin (FICI = 0.31 ± 0.03), bovine serum albumin (BSA, FICI = 0.15 ± 0.17), DNA (FICI = 0.30 ± 0.01), and KCl (FICI = 0.31 ± 0.13) synergized with cefuroxime against USA300 (Supplementary Fig. 8A-D), suggesting the β-lactam sensitization of the USA300 MRSA strain in ASM was triggered by these specific components. No changes were observed when NaCl (FICI = 1.33 ± 0.33), DTPA (FICI = 2.00 ± 0.00), egg yolk emulsion (FICI = 1.58 ± 0.42), or casamino acids (FICI = 1.33 ± 0.33) were introduced to MHB (Supplementary Fig. 8E-H). However, altering the concentrations of mucin, BSA, DNA, and KCl individually to 1/4x and 4x their normal concentration in ASM did not result in any noticeable alteration in cefuroxime MIC in the modified ASM (Supplementary Fig. 8I-L), suggesting potential redundancy in the influence of these multiple components mediating sensitization of USA300 to β-lactams in ASM.

### Investigating the mechanisms of increased resistance to daptomycin in the CF artificial sputum medium

Next, we sought to investigate the mechanisms underlying the increased resistance of USA300 to daptomycin in ASM. We screened the NTML at 4 μg/mL daptomycin, which is equivalent to 1/4x the MIC of USA300 in ASM (Supplementary Fig. 9A). However, no mutants with significantly suppressed growth at the tested daptomycin concentration were identified (Supplementary Fig. 9B), which may be due to genetic redundancies among the actual determinants or the involvement of multiple compensatory or alternative pathways. Alternatively, the underlying determinants might not be represented in the NTML, which covers ∼90% of non-essential genes in USA300, or the phenotype might be due to other non-protein-based mechanisms.

Next, we tested the MIC of daptomycin in SCFM-2 to investigate whether increased resistance would be observed in the other CF sputum-like medium. Interestingly, this phenotype was not maintained in the other CF-mimetic medium, where the daptomycin MIC in SCFM-2 shifted 4-fold below that in MHB, in contrast to the increased resistance in ASM (Supplementary Fig. 9C). Such discrepancy may be due to differences in the media composition, such as the concentration of calcium, required for the activity of daptomycin, where ASM contains DTPA, a chelator, with no calcium added while SCFM-2 contains calcium, or that of other macromolecules (e.g., mucin content is 4-fold different between both media). To this end, we sought to investigate which ASM component may be responsible for altering *S. aureus* susceptibility to daptomycin. As before, we titrated individual ASM components into MHB while determining the daptomycin MIC in checkerboard format. Mucin antagonized with daptomycin against USA300 (Supplementary Fig. 9D, FICI = 4.42 ± 0.08). No changes were observed when BSA (FICI = 2.18 ±0.16), DNA (FICI = 2.01 ± 0.01), KCl (FICI = 2.00 ± 0.00), NaCl (FICI = 2.00 ± 0.00), DTPA (FICI = 2.00 ± 0.00), egg yolk emulsion (FICI = 2.00 ± 0.00), or casamino acids (FICI = 2.17 ± 0.17) were introduced to MHB (Supplementary Fig. 9E-K). Additionally, altering mucin concentrations to 1/4x and 4x its normal ASM concentration produced a dose-dependent increase in MIC with increasing mucin concentrations (Supplementary Fig. 9L). Together, this suggests that the increased resistance to daptomycin in ASM is primarily due to mucin.

Mucin has been shown to bind weakly to daptomycin^59^, thus presenting a potential non-protein-based mechanism for increased daptomycin resistance in ASM. To assess the potential binding, we used well diffusion-based antibiotic bioassays of daptomycin previously exposed to variable concentrations of mucin, revealing a small, but significant, reduction in daptomycin activity in the presence of mucin (Supplementary Fig. 10). Together, our data suggest that the apparent resistance to daptomycin in ASM is mediated by mucin binding.

## Conclusions

Standard AST assays rely on standard lab media, such as MHB, and are used worldwide to guide antimicrobial therapy and study antimicrobial resistance. However, these standardized conditions do not adequately reflect the site of infection, with clinical outcomes often failing to correlate with AST results^13, 14^. Additionally, standard AST conditions have been used for antibiotic discovery, in part as a regulatory requirement to demonstrate efficacy in them for approval^60^. This has likely limited the suite of potential targets and bioactives. There are emerging efforts to study bacterial responses and screen for novel antimicrobials under infection-mimetic conditions to bridge the gap between *in vitro* predictions and *in vivo* outcomes. For example, a screen of FDA-approved drugs identified antimicrobial activity against *Pseudomonas aeruginosa* in CF-sputum-like medium but lower or no activity in the standard MHB, offering a potential for drug repurposing^61^. In this study, we investigated the antimicrobial activity of several antibiotics in CF sputum-mimetic media and found that *S. aureus* was unexpectedly significantly more susceptible to β-lactams and significantly more resistant to daptomycin in CF media than in MHB. We then sought to elucidate the mechanisms underlying these alterations in antibiotic susceptibility.

When investigating genetic determinants of reduced resistance to β-lactams, we uncovered a number of alterations in USA300 that occur under CF sputum-mimetic conditions. These alterations can primarily be grouped by changes in: 1) cell envelope architecture and 2) cellular response mechanisms, especially with regard to cell envelope stress responses. We unveiled remodelling of the MRSA USA300 cell envelope in CF sputum-mimetic conditions, including decreased abundance of PBP2, 2a, 3, and 4 and WTA, decreased polymer length for WTA and LTA, increased cell surface hydrophobicity and a lack of antibiotic-mediated induction of GraXRS-VraFG and stringent response (Fig. 7A). We further characterized alterations to the cell surface of USA300 grown in CF sputum-like media via AFM, which identified alterations to cell surface characteristics that included an increase in surface roughness and cell diameter and a decrease in adhesion between the negatively charged silicon nitride AFM tip and the cell surface, suggesting a relative increase in the overall surface negative charge, in SCFM-2 compared to MHB. The increased susceptibility to cefuroxime in CF sputum-mimetic media was influenced by mucin, BSA, DNA, and KCl in the artificial sputum. Culturing in artificial sputum may induce metabolic reprogramming, influencing β-lactam resistance. For example, glucose-mediated metabolic reprogramming via the pentose phosphate pathway has been shown to increase susceptibility to β-lactams through remodelling of the cell envelope^62^.

**Figure 7.**
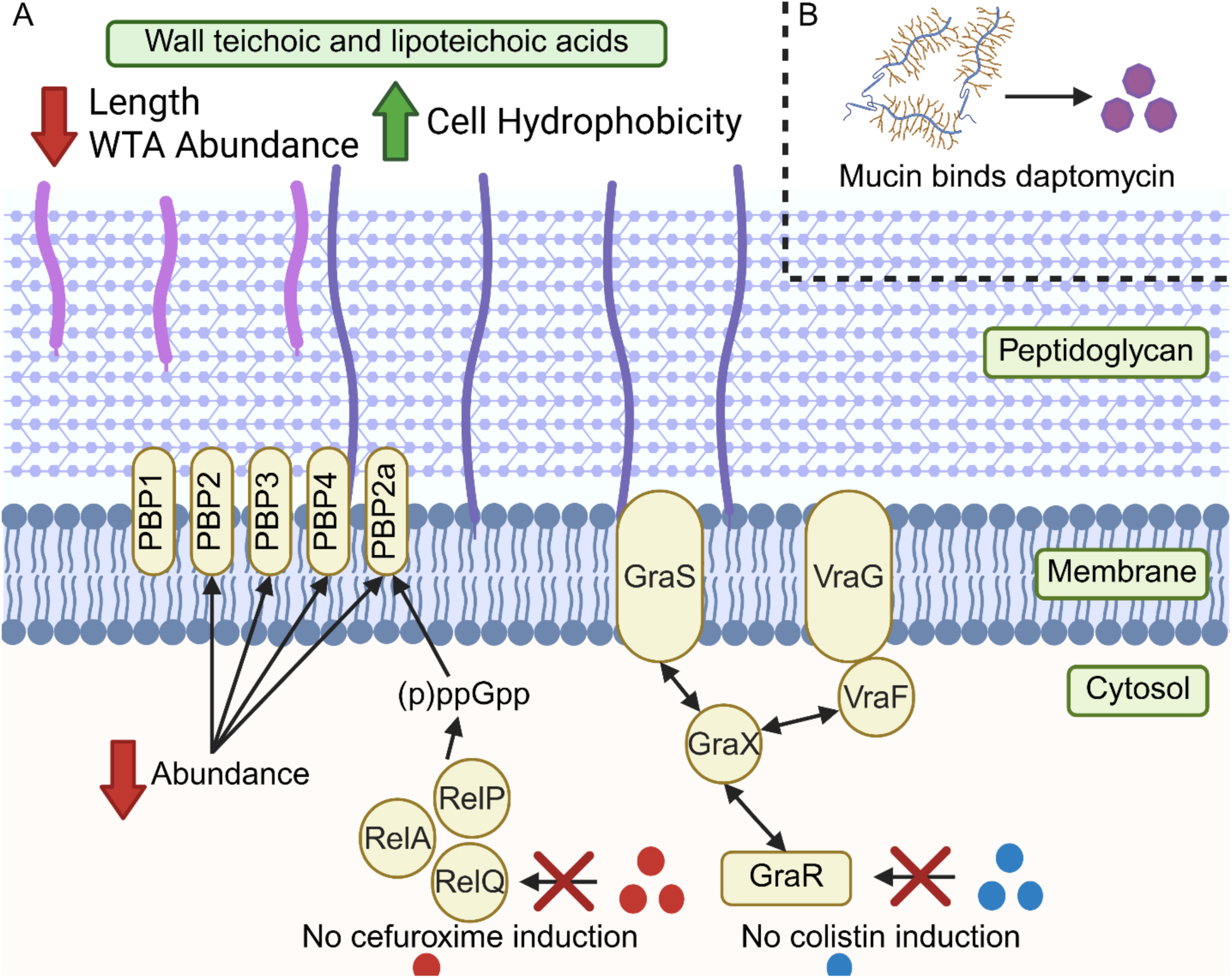
Summary of the current working model for changes underlying the altered susceptibility of MRSA USA300 to β-lactams and daptomycin under CF sputum-mimetic conditions. A) Summary of alterations to USA300 that underlie the increased susceptibility to β-lactams, depicting decreased abundance of PBP2, 2a, 3, and 4 and WTA, decreased polymer length for WTA and LTA, increased cell surface hydrophobicity, and a lack of induction of GraXRS-VraFG and stringent response by colistin and cefuroxime, respectively. B) The apparent daptomycin resistance in ASM is mediated by mucin binding to daptomycin, sequestering it and preventing its action. Created using Biorender.

On the other hand, we identified physicochemical interactions that contribute to the apparent increase in daptomycin resistance of *S. aureus* in ASM. Specifically, mucin directly binds daptomycin, preventing its antimicrobial activity and thereby increasing its effective MIC in ASM (Fig. 7B). Of note, mucin content in ASM is 4-fold higher than in SCFM-2, which explains the opposite phenotype in both media (Supplementary Fig. 9C).

Overall, uncovering antimicrobial susceptibility profiles discordant between infection-mimetic media, such as ASM or SCFM-2, and the standard MHB may enhance our understanding of antibiotic therapeutic responses during infection. Our AST results in CF sputum-mimetic conditions provide insights into bacterial responses under these conditions, uncover mechanistic details underlying the unexpected sensitivity to β-lactams, and could guide and optimize the use of currently available antibiotics. Importantly, our results suggest that β-lactams may be effective in treating MRSA infections in CF patients, warranting further investigation in relevant *in vivo* systems.

## Methods

### Strains, reagents, and culture conditions

Supplementary Table 1 provides details on the strains used in this study. Bacteria were grown in cation-adjusted MHB (herein referred to as MHB), ASM, or SCFM-2 at 37°C, unless otherwise stated. ASM and SCFM-2 were prepared as previously described^19, 63^. The chemical composition of MHB, ASM, and SCFM-2 is detailed in Supplementary Table 2. Due to the opacity of ASM and SCFM-2, assessing bacterial growth turbidimetrically was not feasible. Instead, we used resazurin reduction assays^64^ (Details in Supplementary methods) to indirectly quantify bacterial growth. Overnight cultures used in experiments were prepared in 5 mL tryptic soy broth (TSB) containing relevant selection.

### Isolation of bacteria from liquid cultures

Bacteria were separated from large mucin aggregates present in SCFM-2 liquid cultures by repeated centrifugation at low spin speeds (Supplementary Methods). ASM contains 4x more mucin than SCFM-2, which was harder to completely eliminate using this method. Thus, for assays requiring cell isolation at mid-log phase (i.e., at relatively lower cell density), SCFM-2 was used. Otherwise, ASM was used.

### Antimicrobial susceptibility testing

All MIC assays were conducted according to the Clinical and Laboratory Standards Institute (CLSI) method for MIC testing by broth microdilution^11^ using the colony resuspension method and inoculum size at an OD_600_ of 0.001 in a total volume of 100 µL/well. Plates (96W) were placed in a shaking incubator at 37°C (600 RPM); growth of the bacterial culture was determined by either a resazurin reduction assay (Supplementary Methods) or OD_600_ readings at 24 hours. The fractional inhibitory concentration indices (FICI) were calculated as FICI = (MIC_A_ in combination/MIC_A_ alone) + (MIC_B_ in combination/MIC_B_ alone). FICI values ≤0.5 were interpreted as synergy, >0.5 to <4.0 as additive or indifferent, and >4.0 as antagonistic.

### Nebraska transposon mutant library screen

Overnight cultures of the NTML were prepared by pinning the library into 384-well plates containing LB with 5 μg/mL erythromycin (Sigma-Aldrich) and incubated at 37 °C with shaking (600 rpm) for 24 hours^50^. Subsequently, the overnight culture of the library was used to inoculate clear (MHB) or black (ASM) 384-well plates containing the desired antibiotic concentrations, along with a growth control, using the BioMatrix BM6-BC. Primary screen potential determinants were re-arrayed into 96-well plates in TSB containing 5 μg/mL erythromycin using the BM6-BC and inoculated by pinning from an overnight culture into either clear (MHB) or black (ASM) 96-well plates containing a dilution series of the desired antibiotic using the BM6-BC for follow-up dose response assays. The plates were incubated at 37°C with shaking (600 rpm) for 24 hours, then growth was measured by OD600 for clear plates and by resazurin reduction for black plates. The genomic screen data were normalized in Excel (2021) as previously reported^65^, except that normalization by individual rows and then columns across all plates was performed as the second pass, replacing normalization by well position.

### Membrane protein isolation

Overnight cultures were diluted to an OD_600_ of 0.05 into 60 mL of fresh MHB and SCFM-2 and grown to mid-log phase (OD_600_ of ∼0.8). Cultures were pelleted, washed twice with 5 mL PBS, resuspended in 1 mL PBS and lysed using Lysing Matrix B (MP Biomedicals) in a FastPrep-24™ bead beater. Cell debris was pelleted by centrifugation at 10000xg for 30 minutes at 4°C. Proteins were pelleted via centrifugation at 40000 RPM for 1 hour and resuspended in 50 µL PBS. Membrane protein concentration was determined using the Bradford method with protein assay dye reagent concentrate (Bio-Rad).

### Penicillin binding protein (PBP) quantifications

PBPs were quantified using Bocillin FL (Fisher Scientific) as previously described^66^, with some modifications. PBP2a was quantified using Western blot. Details are provided in Supplementary Methods.

### Lysostaphin-mediated lysis turbidimetric assay

These assays were performed as previously described^53^ using cultures grown in MHB or SCFM-2 (Details in Supplementary Methods).

### Teichoic acid isolation and visualization

Wall teichoic acid (WTA) and lipoteichoic acid (LTA) were isolated from cultures grown overnight on solid MHB and ASM (0.75% agarose) plates using previously established methods^67^, with the following modifications. For WTA isolation, a shaking incubator set to 1000 RPM was used instead of a shaking heat block. For LTA isolation, cells were vortexed for 45 minutes at 4°C in Eppendorf tubes containing 150-212 μm glass beads (Sigma-Aldrich) instead of the MagNA-Lyser. Isolations were separated on a tricine-acrylamide gel as described and stained with 1 mg/mL alcian blue solution. Gels were imaged using a Gel Doc™ EZ Imager (Bio-Rad). The relative abundance of WTA and LTA was determined using imageJ.

### Cell surface hydrophobicity assay

Cell surface hydrophobicity was determined using the microbial adherence to hydrocarbon (MATH) method as previously described^45^.

### Cytochrome C binding assay

The cytochrome C binding assay was performed as previously described^53^ using cells harvested from solid MHB or ASM cultures at an OD_600_ of 15 (Details in Supplementary Methods).

### Bioluminescence promoter-reporter assay

The luciferase expression assays were done as previously described^68^, with some modifications (Supplementary Methods). **(P)ppGpp quantification.** PyDPA was synthesized and verified using previously established procedures^57, 69^. (p)ppGpp was quantified using PyDPA as previously described^70^, with some modifications (Details in Supplementary Methods).

### Atomic force microscopy (AFM) imaging

AFM (Nanowizard 4; JPK Instruments, Berlin, Germany) in quantitative imaging (QI™) mode^71, 72^ was used to analyze cell surface topography and mechanics of USA300 cells cultured in MHB and SCFM-2. Cells were deposited onto clean, Cell-Tak^TM^ (Corning®)-coated coverslips^73^, and imaged using non-conductive silicon nitride cantilevers (Multi-cantilever Tip (MLCT), D cantilever, nominal k = 0.03 N/m and tip radius 20 nm; Bruker, USA). Cantilever spring constant (0.028 N/m) was calibrated using the thermal noise method^74^. QI™ force curves at each pixel of a 128×128 raster scan were collected using a Z-length of 1000 nm, setpoint of 2 nN and a raster scan of 100 µm/s, along with a subset of force curves within a 0.4×0.4 nm square in the center of the cell. Force curves were batch processed using JPK SPM Data Processing software (version 5.1, JPK, Berlin, Germany), and histogram data exported to Excel.

### Antibiotic bioassay by agar well diffusion

Agar well diffusion assays were done following procedures adapted from previously established disc diffusion techniques^75^ as detailed in the Supplementary Methods.

### Statistical analyses

All statistical analyses were calculated using GraphPad Prism 9.

## Acknowledgements

This work was funded by a Saskatchewan Health Research Foundation establishment grant (6115), a Cystic Fibrosis Canada Early Career Investigator Award (1191374), Canada Foundation for Innovation and Innovation Saskatchewan grants, and start-up funds from the Faculty of Science at the University of Regina to O.M.E. O.M.E. holds a Canada Research Chair in Chemogenomics and Antimicrobial Research (CRC-2024-00304). T.J.H. was supported by a Canadian Institutes of Health Research Canada Graduate Scholarship – Master’s (CGS-M) award. Work in the T.E.S.D. laboratory was supported by Natural Science and Engineering Research Council (NSERC DG 2024-06684) and Canada Foundation for Innovation grants to T.E.S.D. The Faculty of Graduate Studies and Research at the University of Regina provided partial support to A.M. This research was conducted on the traditional territories of the Nêhiyawak, Anihsinapek, Nakoda, Dakota, and Lakota, peoples and the homeland of the Métis Nation.

## References

(1) Ciofu, O.; Hansen, C. R.; Hoiby, N. Respiratory bacterial infections in cystic fibrosis. Curr Opin Pulm Med 2013, 19 (3), 251–258. DOI: 10.1097/MCP.0b013e32835f1afc.

(2) Goetz, D. M.; Savant, A. P. Review of CFTR modulators 2020. Pediatric Pulmonology 2021, 56 (12), 3595–3606. DOI: 10.1002/ppul.25627.

(3) Tice, J. A.; Kuntz, K. M.; Wherry, K.; Seidner, M.; Rind, D. M.; Pearson, S. D. The effectiveness and value of novel treatments for cystic fibrosis. Journal of Managed Care & Specialty Pharmacy 2021, 27 (2), 276–280. DOI: 10.18553/jmcp.2021.27.2.276.

(4) Neff, S. L.; Hampton, T. H.; Puerner, C.; Cengher, L.; Doing, G.; Lee, A. J.; Koeppen, K.; Cheung, A. L.; Hogan, D. A.; Cramer, R. A.;, et al. CF-Seq, an accessible web application for rapid re-analysis of cystic fibrosis pathogen RNA sequencing studies. Scientific Data 2022, 9 (1). DOI: 10.1038/s41597-022-01431-1.

(5) Blanchard, A. C.; Waters, V. J. Microbiology of Cystic Fibrosis Airway Disease. Semin Respir Crit Care Med 2019, 40 (6), 727–736. DOI: 10.1055/s-0039-1698464.

(6) Canada, C. F. 2024 Annual Data Report: The Canadian Cystic Fibrosis Registry. Cystic Fibrosis Canada Toronto, Canada: 2025.

(7) Sati, H.; Carrara, E.; Savoldi, A.; Hansen, P.; Garlasco, J.; Campagnaro, E.; Boccia, S.; Castillo-Polo, J. A.; Magrini, E.; Garcia-Vello, P. The WHO Bacterial Priority Pathogens List 2024: a prioritisation study to guide research, development, and public health strategies against antimicrobial resistance. The Lancet infectious diseases 2025.

(8) Wong, J. K.; Ranganathan, S. C.; Hart, E. *Staphylococcus aureus* in early cystic fibrosis lung disease. Pediatric Pulmonology 2013, 48 (12), 1151–1159. DOI: 10.1002/ppul.22863.

(9) Dasenbrook, E. C. Association Between Respiratory Tract Methicillin-Resistant *Staphylococcus aureus* and Survival in Cystic Fibrosis. JAMA 2010, 303 (23), 2386. DOI: 10.1001/jama.2010.791.

(10) Ren, C. L.; Morgan, W. J.; Konstan, M. W.; Schechter, M. S.; Wagener, J. S.; Fisher, K. A.; Regelmann, W. E. Presence of methicillin resistant *Staphylococcus aureus* in respiratory cultures from cystic fibrosis patients is associated with lower lung function. Pediatric Pulmonology 2007, 42 (6), 513–518. DOI: 10.1002/ppul.20604.

(11) CLSI. Methods for Dilution Antimicrobial Susceptibility Tests for Bacteria That Grow Aerobically; Approved Standard—Ninth Edition. CLSI document M07-A9.; Clinical and Laboratory Standards Institute, 2012.

(12) Mueller, J. H.; Hinton, J. A protein-free medium for primary isolation of the Gonococcus and Meningococcus. Proc. Soc. Expt. Biol. Med. 1941, 48 (1), 330–333. DOI: 10.3181/00379727-48-13311.

(13) Hurley, M. N.; Ariff, A. H.; Bertenshaw, C.; Bhatt, J.; Smyth, A. R. Results of antibiotic susceptibility testing do not influence clinical outcome in children with cystic fibrosis. J Cyst Fibros 2012, 11 (4), 288–292. DOI: 10.1016/j.jcf.2012.02.006.

(14) Smith, A. L.; Fiel, S. B.; Mayer-Hamblett, N.; Ramsey, B.; Burns, J. L. Susceptibility testing of *Pseudomonas aeruginosa* isolates and clinical response to parenteral antibiotic administration: lack of association in cystic fibrosis. Chest 2003, 123 (5), 1495–1502.

(15) Macgowan, A. P.; Surveillance, B. W. P. o. R. Clinical implications of antimicrobial resistance for therapy. J Antimicrob Chemother 2008, 62 Suppl 2, ii105–114. DOI: 10.1093/jac/dkn357.

(16) Tong, M.; French, S.; El Zahed, S. S.; Ong, W. K.; Karp, P. D.; Brown, E. D. Gene Dispensability in *Escherichia coli* Grown in Thirty Different Carbon Environments. mBio 2020, 11 (5). DOI: 10.1128/mBio.02259-20.

(17) Fung, C.; Naughton, S.; Turnbull, L.; Tingpej, P.; Rose, B.; Arthur, J.; Hu, H.; Harmer, C.; Harbour, C.; Hassett, D. J.;, et al. Gene expression of *Pseudomonas aeruginosa* in a mucin-containing synthetic growth medium mimicking cystic fibrosis lung sputum. J Med Microbiol 2010, 59 (Pt 9), 1089–1100. DOI: 10.1099/jmm.0.019984-0.

(18) Quinn, R. A.; Whiteson, K.; Lim, Y. W.; Salamon, P.; Bailey, B.; Mienardi, S.; Sanchez, S. E.; Blake, D.; Conrad, D.; Rohwer, F. A Winogradsky-based culture system shows an association between microbial fermentation and cystic fibrosis exacerbation. ISME J 2015, 9 (4), 1024–1038. DOI: 10.1038/ismej.2014.234.

(19) Turner, K. H.; Wessel, A. K.; Palmer, G. C.; Murray, J. L.; Whiteley, M. Essential genome of *Pseudomonas aeruginosa* in cystic fibrosis sputum. Proc Natl Acad Sci U S A 2015, 112 (13), 4110–4115. DOI: 10.1073/pnas.1419677112.

(20) Diaz Iglesias, Y.; Wilms, T.; Vanbever, R.; Van Bambeke, F. Activity of Antibiotics against *Staphylococcus aureus* in an *In Vitro* Model of Biofilms in the Context of Cystic Fibrosis: Influence of the Culture Medium. Antimicrob Agents Chemother 2019, 63 (7). DOI: 10.1128/AAC.00602-19.

(21) Tenover, F. C.; Goering, R. V. Methicillin-resistant Staphylococcus aureus strain USA300: origin and epidemiology. Journal of Antimicrobial Chemotherapy 2009, 64 (3), 441–446.

(22) Andrade, F. F.; Silva, D.; Rodrigues, A.; Pina-Vaz, C. Colistin update on its mechanism of action and resistance, present and future challenges. Microorganisms 2020, 8 (11), 1716.

(23) Bycroft, B. W.; Shute, R. E. The Molecular Basis for the Mode of Action of Beta-Lactam Antibiotics and Mechanisms of Resistance. Pharmaceutical Research 1985, 02 (1), 3–14. DOI: 10.1023/a:1016305704057.

(24) Gray, D. A.; Wenzel, M. More Than a Pore: A Current Perspective on the In Vivo Mode of Action of the Lipopeptide Antibiotic Daptomycin. Antibiotics 2020, 9 (1), 17. DOI: 10.3390/antibiotics9010017.

(25) Osorio, C.; Garzón, L.; Jaimes, D.; Silva, E.; Bustos, R.-H. Impact on Antibiotic Resistance, Therapeutic Success, and Control of Side Effects in Therapeutic Drug Monitoring (TDM) of Daptomycin: A Scoping Review. Antibiotics 2021, 10 (3), 263. DOI: 10.3390/antibiotics10030263.

(26) Silverman, J. A.; Mortin, L. I.; VanPraagh, A. D. G.; Li, T.; Alder, J. Inhibition of Daptomycin by Pulmonary Surfactant: *In Vitro* Modeling and Clinical Impact. The Journal of Infectious Diseases 2005, 191 (12), 2149–2152. DOI: 10.1086/430352 (accessed 1/27/2026).

(27) Harada, Y.; Yanagihara, K.; Yamada, K.; Migiyama, Y.; Nagaoka, K.; Morinaga, Y.; Nakamura, S.; Imamura, Y.; Hasegawa, H.; Miyazaki, T.;, et al. *In vivo* efficacy of daptomycin against methicillin-resistant *Staphylococcus aureus* in a mouse model of hematogenous pulmonary infection. Antimicrob Agents Chemother 2013, 57 (6), 2841–2844. DOI: 10.1128/aac.02331-12 From NLM.

(28) Henken, S.; Bohling, J.; Martens-Lobenhoffer, J.; Paton, J. C.; Ogunniyi, A. D.; Briles, D. E.; Salisbury, V. C.; Wedekind, D.; Bode-Böger, S. M.; Welsh, T.;, et al. Efficacy Profiles of Daptomycin for Treatment of Invasive and Noninvasive Pulmonary Infections with *Streptococcus pneumoniae*. Antimicrob Agents Ch 2010, 54 (2), 707–717. DOI: doi:10.1128/aac.00943-09.

(29) CLSI. Performance Standards for Antimicrobial Susceptibility Testing CLSI supplement M100. Approved Standard, Pennsylvania 2023, 33rd ed.

(30) Christianson, S.; Golding, G. R.; Campbell, J.; Mulvey, M. R. Comparative genomics of Canadian epidemic lineages of methicillin-resistant Staphylococcus aureus. Journal of Clinical Microbiology 2007, 45 (6), 1904–1911.

(31) Fey, P. D.; Endres, J. L.; Yajjala, V. K.; Widhelm, T. J.; Boissy, R. J.; Bose, J. L.; Bayles, K. W. A genetic resource for rapid and comprehensive phenotype screening of nonessential *Staphylococcus aureus* genes. MBio 2013, 4 (1), e00537–00512. DOI: 10.1128/mBio.00537-12.

(32) Abdelmalek, N.; Yousief, S. W.; Tanca, A.; Bojer, M. S.; Alobaidallah, M. S. A.; Olsen, J. E.; Paglietti, B. Exploring the susceptome of methicillin-resistant Staphylococcus aureus to oxacillin and cefazolin using integrated TraDIS and proteomics. International Journal of Antimicrobial Agents 2025, 107652.

(33) Mlynek, K. D.; Callahan, M. T.; Shimkevitch, A. V.; Farmer, J. T.; Endres, J. L.; Marchand, M.; Bayles, K. W.; Horswill, A. R.; Kaplan, J. B. Effects of low-dose amoxicillin on Staphylococcus aureus USA300 biofilms. Antimicrobial agents and chemotherapy 2016, 60 (5), 2639–2651.

(34) Losey, H. C.; Ruthenburg, A. J.; Verdine, G. L. Crystal structure of Staphylococcus aureus tRNA adenosine deaminase TadA in complex with RNA. Nature structural & molecular biology 2006, 13 (2), 153–159.

(35) Fuchs, S.; Mehlan, H.; Bernhardt, J.; Hennig, A.; Michalik, S.; Surmann, K.; Pané-Farré, J.; Giese, A.; Weiss, S.; Backert, L. AureoWiki ̵ The repository of the Staphylococcus aureus research and annotation community. International journal of medical microbiology 2018, 308 (6), 558–568.

(36) Santa Maria, J. P.; Sadaka, A.; Moussa, S. H.; Brown, S.; Zhang, Y. J.; Rubin, E. J.; Gilmore, M. S.; Walker, S. Compound-gene interaction mapping reveals distinct roles for *Staphylococcus aureus* teichoic acids. Proceedings of the National Academy of Sciences 2014, 111 (34), 12510–12515. DOI: doi:10.1073/pnas.1404099111.

(37) Łeski, T. A.; Tomasz, A. Role of penicillin-binding protein 2 (PBP2) in the antibiotic susceptibility and cell wall cross-linking of Staphylococcus aureus: evidence for the cooperative functioning of PBP2, PBP4, and PBP2A. Journal of bacteriology 2005, 187 (5), 1815–1824.

(38) Bernal, P.; Lemaire, S.; Pinho, M. G.; Mobashery, S.; Hinds, J.; Taylor, P. W. Insertion of epicatechin gallate into the cytoplasmic membrane of methicillin-resistant Staphylococcus aureus disrupts penicillin-binding protein (PBP) 2a-mediated β-lactam resistance by delocalizing PBP2. Journal of Biological Chemistry 2010, 285 (31), 24055–24065.

(39) Göhring, N.; Fedtke, I.; Xia, G.; Jorge, A. M.; Pinho, M. G.; Bertsche, U.; Peschel, A. New role of the disulfide stress effector YjbH in β-lactam susceptibility of Staphylococcus aureus. Antimicrobial agents and chemotherapy 2011, 55 (12), 5452–5458.

(40) Karinou, E.; Schuster, C. F.; Pazos, M.; Vollmer, W.; Gründling, A. Inactivation of the monofunctional peptidoglycan glycosyltransferase SgtB allows Staphylococcus aureus to survive in the absence of lipoteichoic acid. Journal of Bacteriology 2019, 201 (1), 10.1128/jb. 00574–00518.

(41) Sewell, E. W.; Brown, E. D. Taking aim at wall teichoic acid synthesis: new biology and new leads for antibiotics. The Journal of antibiotics 2014, 67 (1), 43–51.

(42) Fedtke, I.; Mader, D.; Kohler, T.; Moll, H.; Nicholson, G.; Biswas, R.; Henseler, K.; Götz, F.; Zähringer, U.; Peschel, A. A Staphylococcus aureus ypfP mutant with strongly reduced lipoteichoic acid (LTA) content: LTA governs bacterial surface properties and autolysin activity. Molecular microbiology 2007, 65 (4), 1078–1091.

(43) Kohler, T.; Weidenmaier, C.; Peschel, A. Wall teichoic acid protects Staphylococcus aureus against antimicrobial fatty acids from human skin. Journal of bacteriology 2009, 191 (13), 4482–4484.

(44) Jeon, H.; Lee, H.; Song, C.; Lee, I. G. Structural Insights into the Staphylococcus aureus DltC-Mediated D-Alanine Transfer. Biomolecules 2025, 16 (1). DOI: 10.3390/biom16010044 From NLM.

(45) Rosenberg, M.; Gutnick, D.; Rosenberg, E. Adherence of bacteria to hydrocarbons: a simple method for measuring cell-surface hydrophobicity. FEMS Microbiol Lett 1980, 9 (1), 29–33.

(46) Hesser, A. R.; Matano, L. M.; Vickery, C. R.; Wood, B. M.; Santiago, A. G.; Morris, H. G.; Do, T.; Losick, R.; Walker, S. The length of lipoteichoic acid polymers controls Staphylococcus aureus cell size and envelope integrity. Journal of bacteriology 2020, 202 (16), 10.1128/jb. 00149–00120.

(47) Gründling, A.; Schneewind, O. Synthesis of glycerol phosphate lipoteichoic acid in Staphylococcus aureus. Proceedings of the National Academy of Sciences 2007, 104 (20), 8478–8483.

(48) Mikkelsen, K.; Sirisarn, W.; Alharbi, O.; Alharbi, M.; Liu, H.; Nøhr-Meldgaard, K.; Mayer, K.; Vestergaard, M.; Gallagher, L. A.; Derrick, J. P. The novel membrane-associated auxiliary factors AuxA and AuxB modulate β-lactam resistance in MRSA by stabilizing lipoteichoic acids. International journal of antimicrobial agents 2021, 57 (3), 106283.

(49) Falord, M.; Karimova, G.; Hiron, A.; Msadek, T. GraXSR proteins interact with the VraFG ABC transporter to form a five-component system required for cationic antimicrobial peptide sensing and resistance in *Staphylococcus aureus*. Antimicrob Agents Chemother 2012, 56 (2), 1047–1058. DOI: 10.1128/AAC.05054-11.

(50) El-Halfawy, O. M.; Czarny, T. L.; Flannagan, R. S.; Day, J.; Bozelli, J. C., Jr.; Kuiack, R. C.; Salim, A.; Eckert, P.; Epand, R. M.; McGavin, M. J.;, et al. Discovery of an antivirulence compound that reverses beta-lactam resistance in MRSA. Nat Chem Biol 2020, 16 (2), 143–149. DOI: 10.1038/s41589-019-0401-8 From NLM Medline.

(51) Meehl, M.; Herbert, S.; Götz, F.; Cheung, A. Interaction of the GraRS two-component system with the VraFG ABC transporter to support vancomycin-intermediate resistance in Staphylococcus aureus. Antimicrobial agents and chemotherapy 2007, 51 (8), 2679–2689.

(52) Yang, S.-J.; Mishra, N. N.; Rubio, A.; Bayer, A. S. Causal role of single nucleotide polymorphisms within the mprF gene of Staphylococcus aureus in daptomycin resistance. Antimicrobial agents and chemotherapy 2013, 57 (11), 5658–5664.

(53) Douglas, E. J. A.; Palk, N.; Brignoli, T.; Altwiley, D.; Boura, M.; Laabei, M.; Recker, M.; Cheung, G. Y. C.; Liu, R.; Hsieh, R. C.;, et al. Extensive re-modelling of the cell wall during the development of *Staphylococcus aureus* bacteraemia. eLife 2023. DOI: 10.7554/elife.87026.2.

(54) Geiger, T.; Kastle, B.; Gratani, F. L.; Goerke, C.; Wolz, C. Two small (p)ppGpp synthases in *Staphylococcus aureus* mediate tolerance against cell envelope stress conditions. J Bacteriol 2014, 196 (4), 894–902. DOI: 10.1128/JB.01201-13 From NLM Medline.

(55) Yang, N.; Xie, S.; Tang, N.-Y.; Choi, M. Y.; Wang, Y.; Watt, R. M. The Ps and Qs of alarmone synthesis in Staphylococcus aureus. PloS one 2019, 14 (10), e0213630.

(56) Mwangi, M. M.; Kim, C.; Chung, M.; Tsai, J.; Vijayadamodar, G.; Benitez, M.; Jarvie, T. P.; Du, L.; Tomasz, A. Whole-genome sequencing reveals a link between β-lactam resistance and synthetases of the alarmone (p) ppGpp in Staphylococcus aureus. Microbial Drug Resistance 2013, 19 (3), 153–159.

(57) Rhee, H.-W.; Lee, C.-R.; Cho, S.-H.; Song, M.-R.; Cashel, M.; Choy, H. E.; Seok, Y.-J.; Hong, J.-I. Selective Fluorescent Chemosensor for the Bacterial Alarmone (p)ppGpp. Journal of the American Chemical Society 2008, 130 (3), 784–785. DOI: 10.1021/ja0759139.

(58) Bhawini, A.; Pandey, P.; Dubey, A. P.; Zehra, A.; Nath, G.; Mishra, M. N. RelQ mediates the expression of β-lactam resistance in methicillin-resistant Staphylococcus aureus. Frontiers in Microbiology 2019, 10, 339.

(59) Huang, J. X.; Blaskovich, M. A.; Pelingon, R.; Ramu, S.; Kavanagh, A.; Elliott, A. G.; Butler, M. S.; Montgomery, A. B.; Cooper, M. A. Mucin Binding Reduces Colistin Antimicrobial Activity. Antimicrob Agents Chemother 2015, 59 (10), 5925–5931. DOI: 10.1128/AAC.00808-15 From NLM Medline.

(60) van Belkum, A.; Bachmann, T. T.; Lüdke, G.; Lisby, J. G.; Kahlmeter, G.; Mohess, A.; Becker, K.; Hays, J. P.; Woodford, N.; Mitsakakis, K. Developmental roadmap for antimicrobial susceptibility testing systems. Nature Reviews Microbiology 2019, 17 (1), 51–62.

(61) Di Bonaventura, G.; Lupetti, V.; Di Giulio, A.; Muzzi, M.; Piccirilli, A.; Cariani, L.; Pompilio, A. Repurposing high-throughput screening identifies unconventional drugs with antibacterial and antibiofilm activities against Pseudomonas aeruginosa under experimental conditions relevant to cystic fibrosis. Microbiology Spectrum 2023, 11 (4), e00352–00323.

(62) Zeden, M. S.; Gallagher, L. A.; Bueno, E.; Nolan, A. C.; Ahn, J.; Shinde, D.; Razvi, F.; Sladek, M.; Burke, Ó.; O’Neill, E. Metabolic reprogramming and altered cell envelope characteristics in a pentose phosphate pathway mutant increases MRSA resistance to β-lactam antibiotics. PLoS pathogens 2023, 19 (7), e1011536.

(63) El-Halfawy, O. M.; Naguib, M. M.; Valvano, M. A. Novel antibiotic combinations proposed for treatment of *Burkholderia cepacia* complex infections. Antimicrobial Resistance & Infection Control 2017, 6 (1). DOI: 10.1186/s13756-017-0279-8.

(64) Riss, T. L.; Moravec, R. A.; Niles, A. L.; Duellman, S.; Benink, H. A.; Worzella, T. J.; Minor, L. Cell viability assays. Assay Guidance Manual [Internet] 2016.

(65) Mangat, C. S.; Bharat, A.; Gehrke, S. S.; Brown, E. D. Rank ordering plate data facilitates data visualization and normalization in high-throughput screening. J Biomol Screen 2014, 19 (9), 1314–1320. DOI: 10.1177/1087057114534298 From NLM Medline.

(66) Kocaoglu, O.; Carlson, E. E. Penicillin-binding protein imaging probes. Current protocols in chemical biology 2013, 5 (4), 239–250.

(67) Kho, K.; Meredith, T. C. Extraction and analysis of bacterial teichoic acids. Bio-protocol 2018, 8 (21), e3078–e3078.

(68) Flannagan, R. S.; Kuiack, R. C.; McGavin, M. J.; Heinrichs, D. E. Staphylococcus aureus uses the GraXRS regulatory system to sense and adapt to the acidified phagolysosome in macrophages. MBio 2018, 9 (4), 10.1128/mbio. 01143–01118.

(69) Conti, G.; Minneci, M.; Sattin, S. Optimised Synthesis of the Bacterial Magic Spot (p)ppGpp Chemosensor PyDPA. ChemBioChem 2019, 20 (13), 1717–1721. DOI: 10.1002/cbic.201900013 (accessed 2026/03/29).

(70) Gao, W.; Chua, K.; Davies, J. K.; Newton, H. J.; Seemann, T.; Harrison, P. F.; Holmes, N. E.; Rhee, H.-W.; Hong, J.-I.; Hartland, E. L. Two novel point mutations in clinical Staphylococcus aureus reduce linezolid susceptibility and switch on the stringent response to promote persistent infection. PLoS pathogens 2010, 6 (6), e1000944.

(71) Chopinet, L.; Formosa, C.; Rols, M.; Duval, R.; Dague, E. Imaging living cells surface and quantifying its properties at high resolution using AFM in QI™ mode. Micron 2013, 48, 26–33.

(72) Bhat, S. V.; Sultana, T.; Körnig, A.; McGrath, S.; Shahina, Z.; Dahms, T. E. Correlative atomic force microscopy quantitative imaging-laser scanning confocal microscopy quantifies the impact of stressors on live cells in real-time. Scientific reports 2018, 8 (1), 8305.

(73) Shahina, Z.; Bhat, S. V.; Ndlovu, E.; Sultana, T.; Körnig, A.; Dague, É.; Dahms, T. E. Cellulomics of Live Yeast by Advanced and Correlative Microscopy. In *Laboratory Protocols in Fungal Biology: Current Methods in Fungal Biology*, Springer, 2022; pp 159–174.

(74) Lübbe, J.; Temmen, M.; Rahe, P.; Kühnle, A.; Reichling, M. Determining cantilever stiffness from thermal noise. Beilstein journal of nanotechnology 2013, 4 (1), 227–233.

(75) Andrews, J. BSAC standardized disc susceptibility testing method (version 8). Journal of antimicrobial chemotherapy 2009, 64 (3), 454–489.

