## Supplementary File for "Clinically-relevant altered antibiotic responses and mechanisms of β-lactam sensitization of MRSA in cystic fibrosis artificial sputum"

**Supplementary Figures, Tables, and Methods**

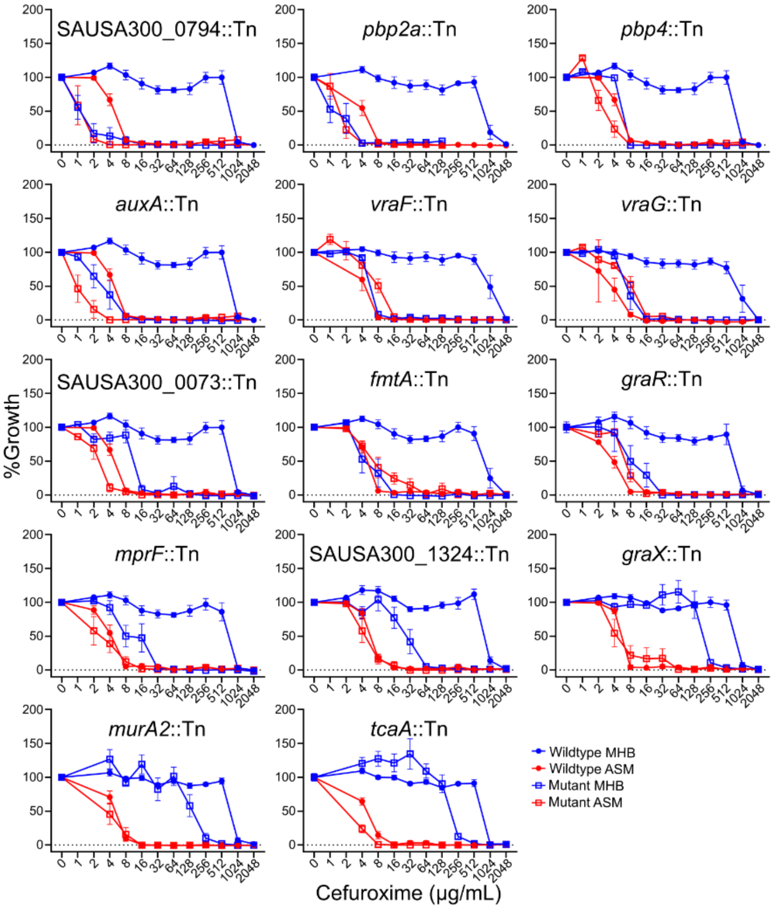


**Supplementary Figure 1. MIC of cefuroxime against NTML mutants of putative determinants of cefuroxime resistance in MHB.** MIC dose-response growth curves for mutants that are more sensitive to cefuroxime in MHB, shown as mean ± SEM, n = 6 from three independent experiments.

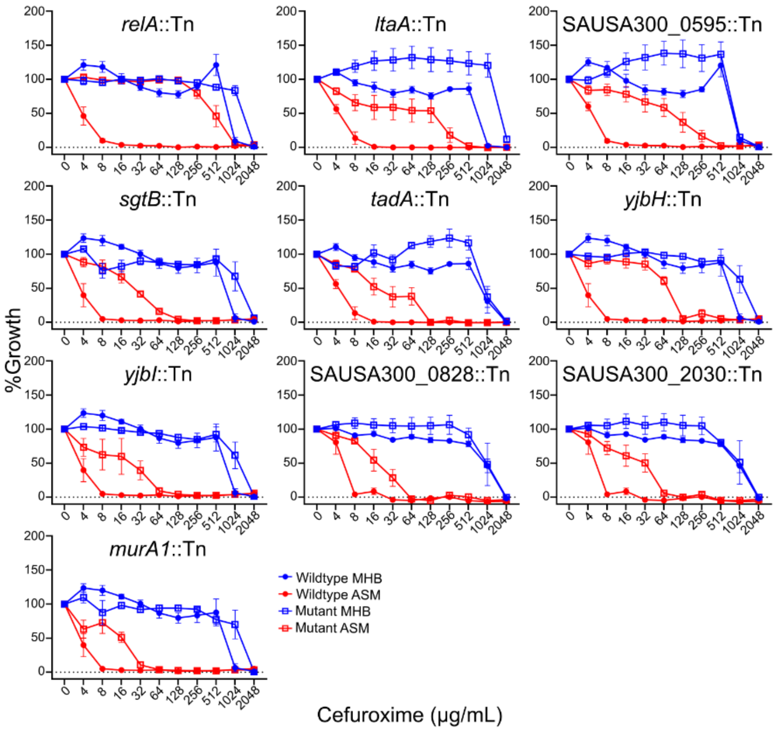


**Supplementary Figure 2. MIC of cefuroxime against NTML mutants of putative determinants of cefuroxime sensitivity in ASM.** MIC dose-response growth curves for mutants that are more resistant to cefuroxime in ASM, shown as mean ± SEM, n = 6 from three independent experiments.

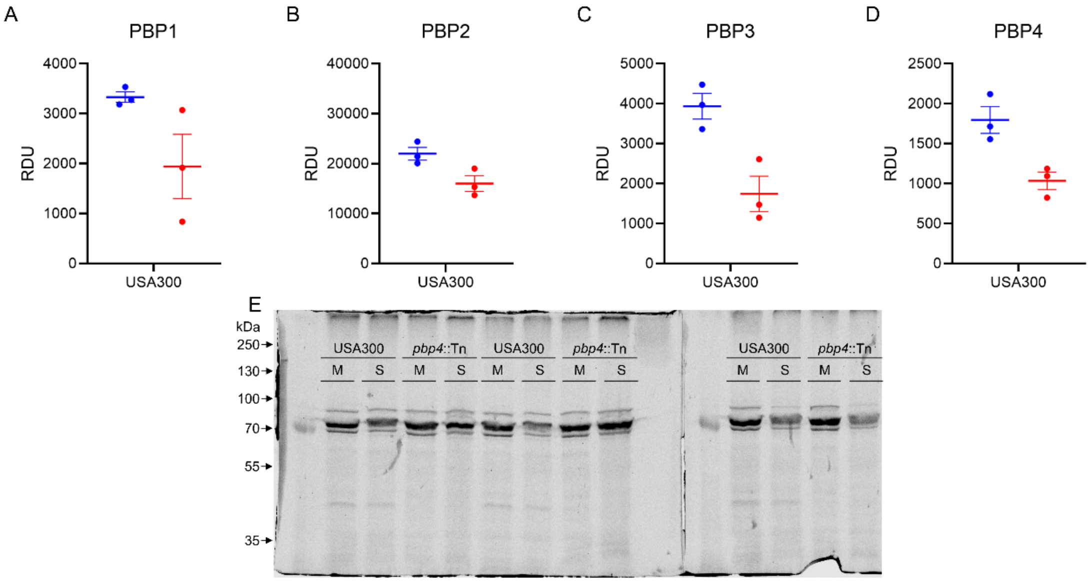


**Supplementary Figure 3. BOCILLIN FL-labelled PBP visualization and quantification.** Values for the density units (RDU) obtained from quantifying A) PBP1, B) PBP2, C) PBP3, and D) PBP4 from cells isolated from MHB (blue) or SCFM-2 (red). E) BOCILLIN FL-labelled gel image for all isolations. M: MHB, S: SCFM-2. Replicates shown are from three independent isolations (n=3).

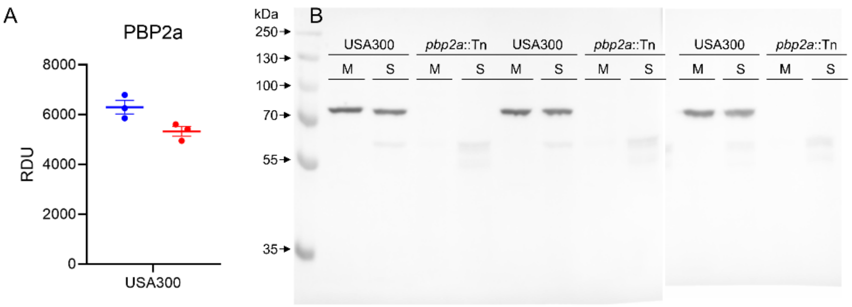


**Supplementary Figure 4. PBP2a Western blot quantification.** A) Values for the density units (RDU) obtained from quantifying PBP2a from cells isolated from MHB (blue) or SCFM-2 (red). B) Western blot image for all isolations. M: MHB, S: SCFM-2. Replicates shown are from three independent isolations (n = 3)

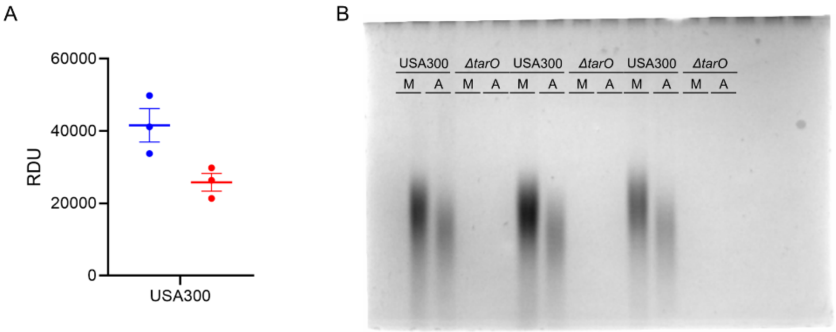


32 **Supplementary Figure 5. WTA visualization and quantification.** A) Values for relative density units (RDU) obtained from quantifying crude WTA isolations from cells isolated from MHB (blue) or ASM (red). B) Gel images for all isolations. M: MHB, A: ASM. Replicates shown are from three independent isolations (n = 3)

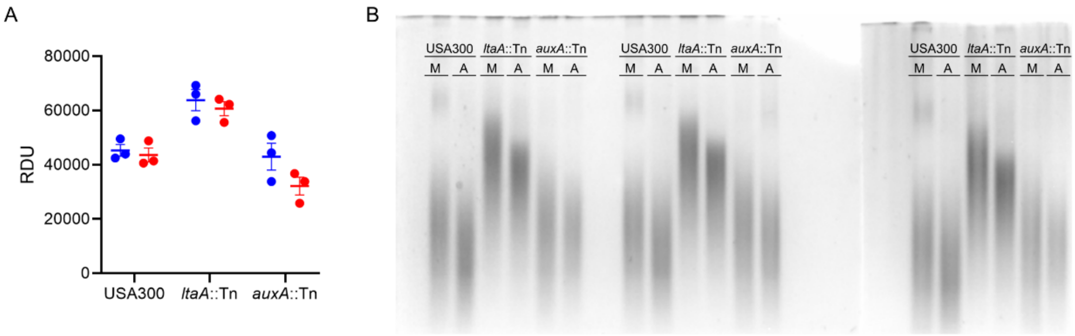


**Supplementary Figure 6. LTA visualization and quantification.** A) Values for relative density units (RDU) obtained from quantifying crude LTA isolations from cells isolated from MHB (blue) or SCFM-2 (red). B) Gel images for all isolations. Replicates shown are from three independent isolations (n = 3)

A

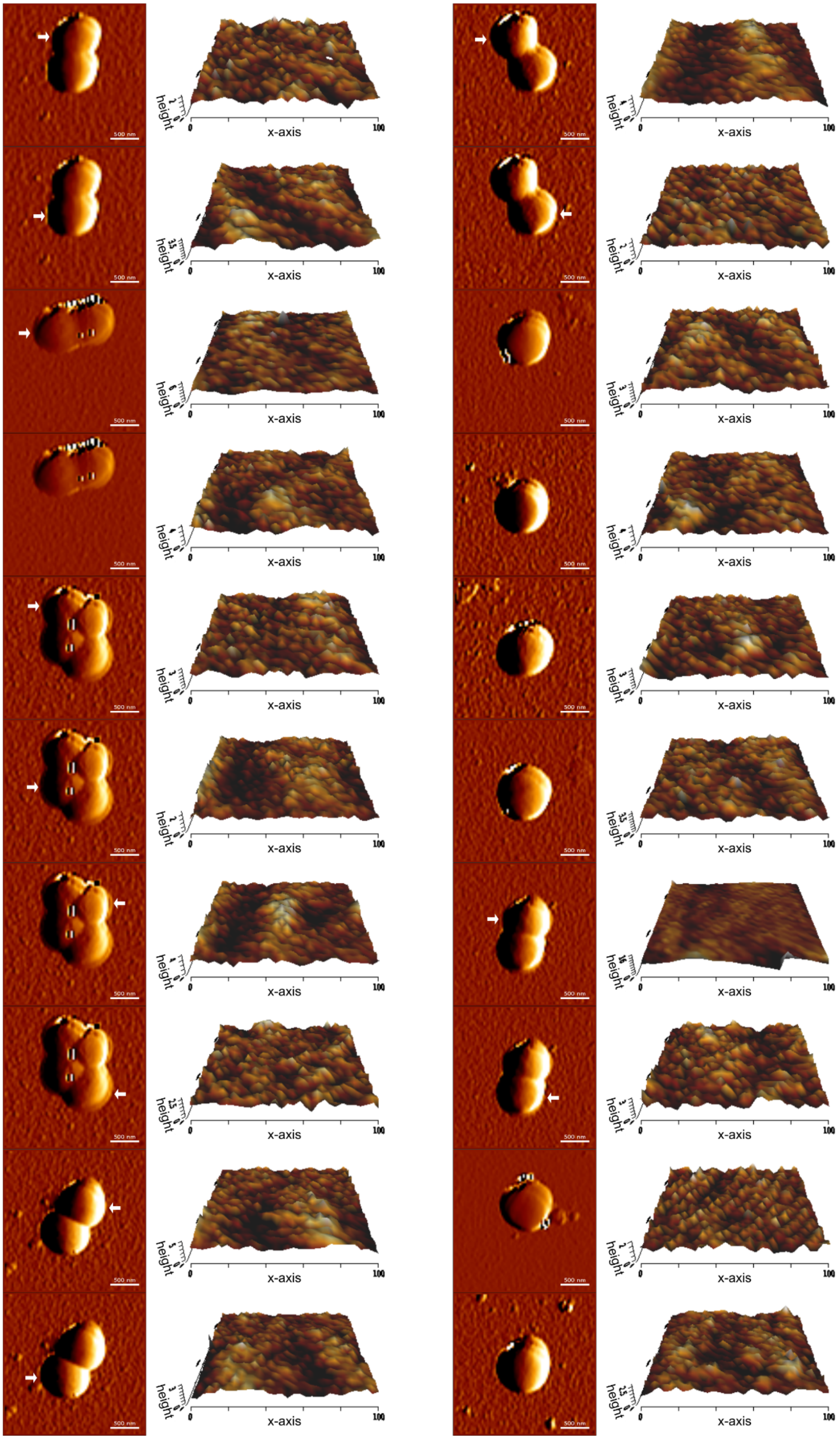

B

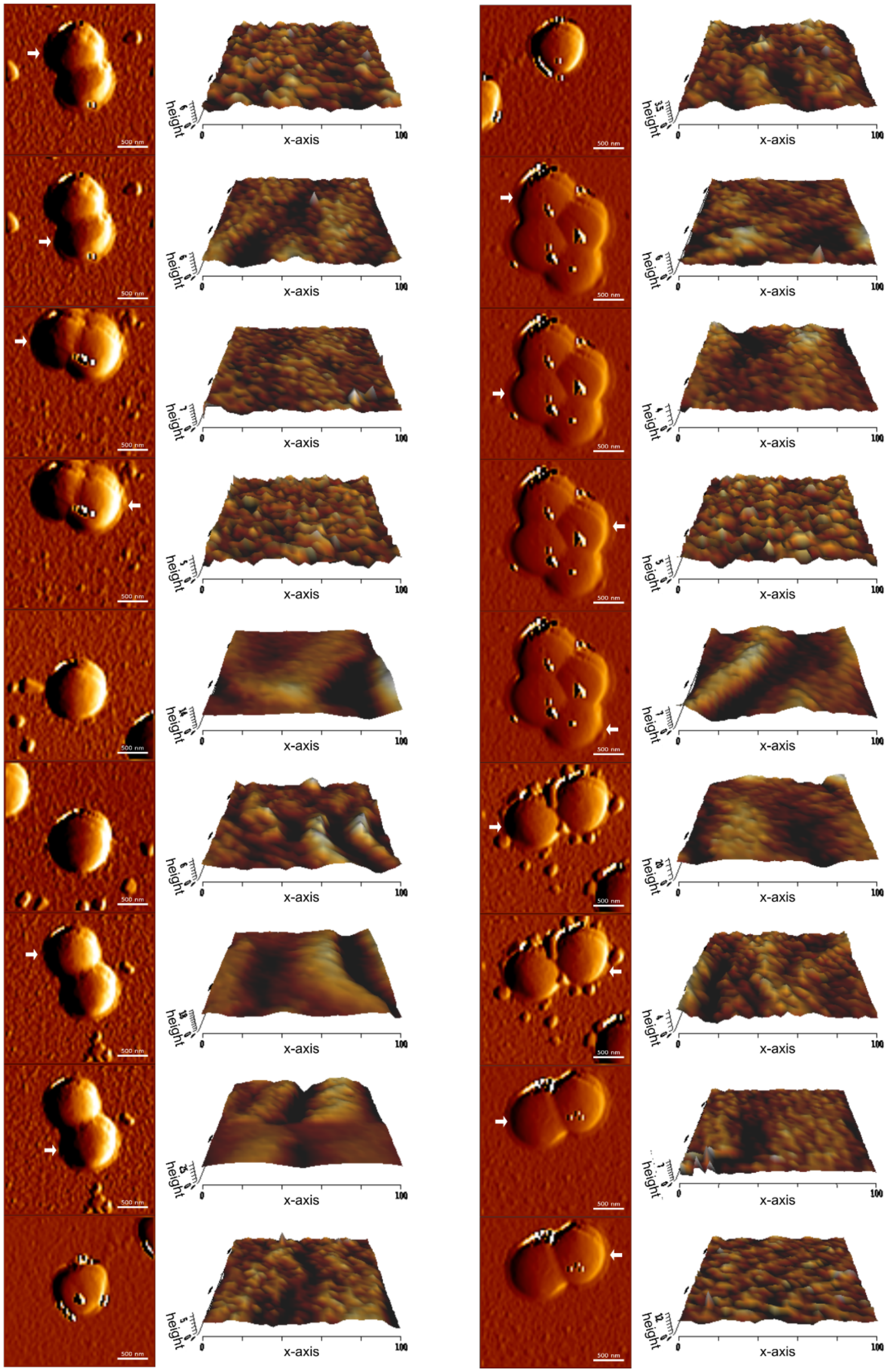


**Supplementary Figure 7. AFM images used for quantification of cell surface properties.** Whole cell images and 3D images of cell topography obtained from AFM imaging of USA300 grown in A) MHB and B) SCFM-2. Images represent a minimum of 18 cells from three biological replicates. In cases where there are multiple cells in the field of view, the cell used in the quantification and whose 3D topography shown is marked with a white arrow.

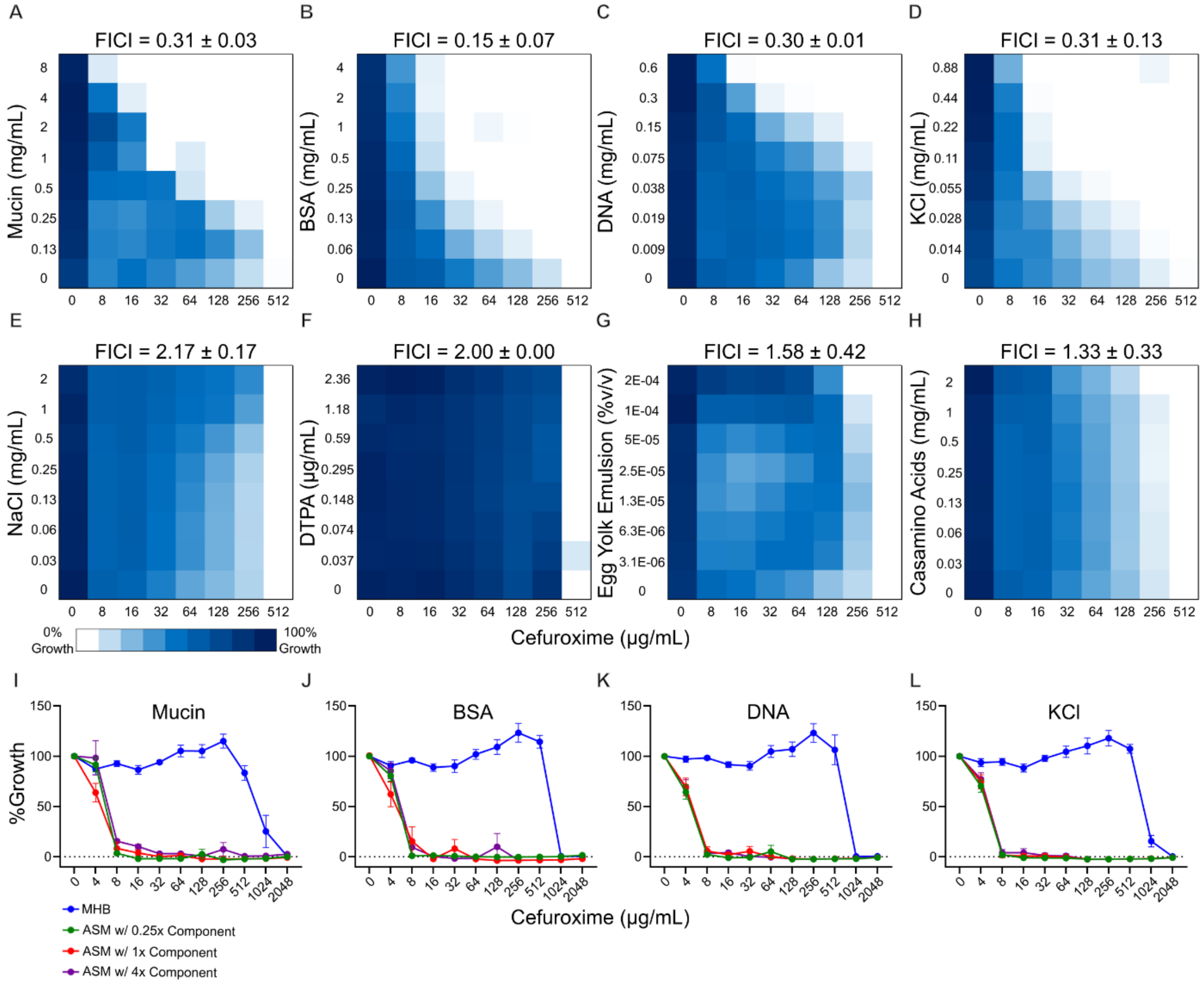


**Supplementary Figure 8. Multiple ASM components influence USA300 susceptibility to cefuroxime.** A-H) Representative checkerboard assays of individual ASM components with cefuroxime in MHB against S. aureus USA300. The highest tested concentration of all ASM ingredients corresponds to 40% of their normal concentrations in ASM. FICI values are reported as mean ± SEM (n = 3). I-L) Cefuroxime MICs performed in ASM with varying concentrations of I) mucin, J) BSA, K) DNA, and L) KCl compared to MHB against USA300, shown as mean ± SEM, n = 6 from three independent experiments.

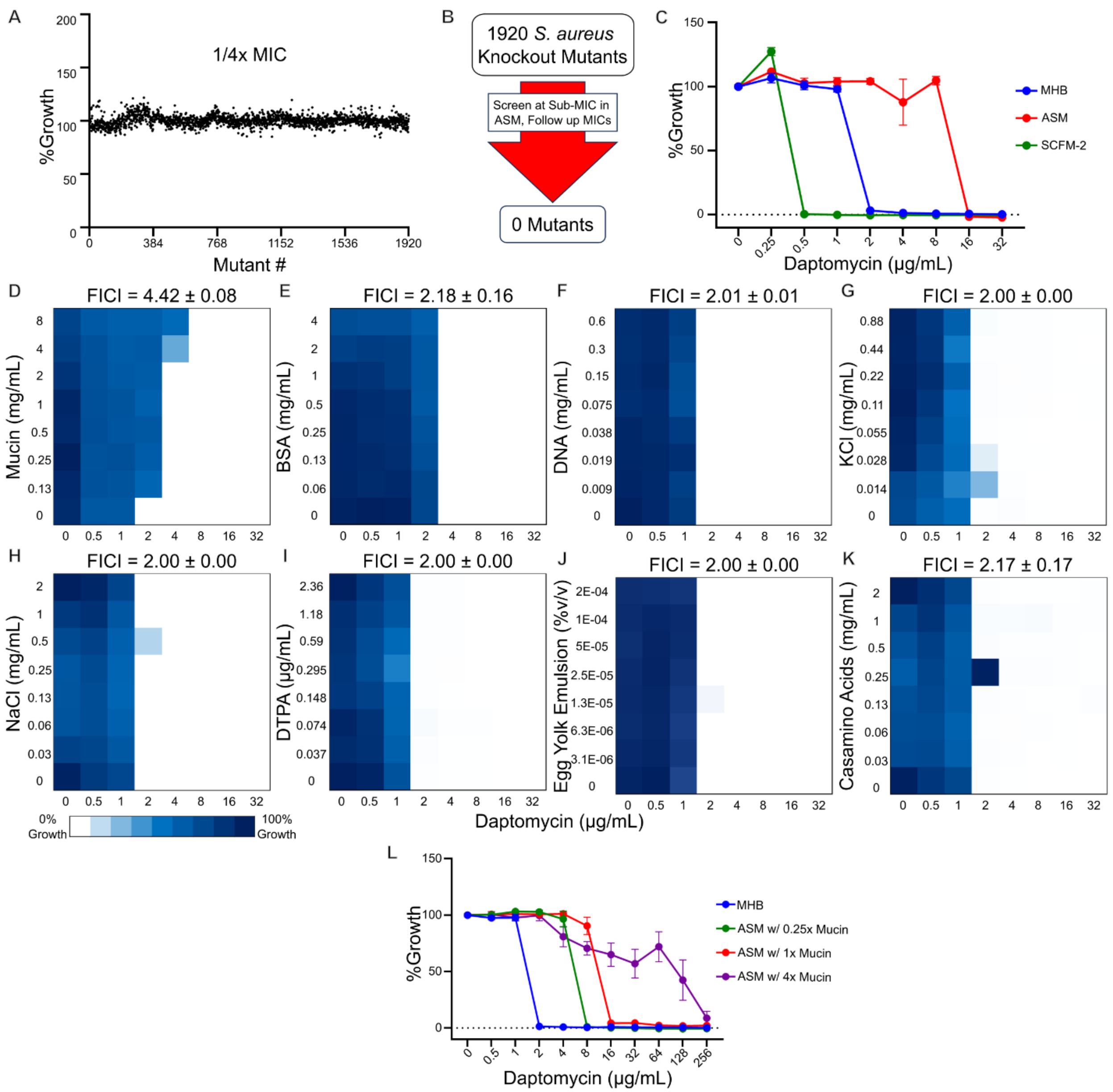


**Supplementary Figure 9. The apparent daptomycin resistance in ASM is likely mediated by mucin binding.** A) Index plot for the NTML screened at ¼ MIC of daptomycin in ASM. B) Workflow of NTML screen follow-up, which did not identify any genetic determinants of daptomycin resistance. C) MIC assay for daptomycin against USA300 performed in MHB, ASM, or SCFM-2, shown as mean ± SEM, n = 6 from three independent experiments. D-K) Representative checkerboard assays performed against USA300 in MHB with daptomycin in combination with D) mucin, E) BSA, F) DNA, G) KCl,

H) NaCl, I) DTPA, J) egg yolk emulsion, and K) casamino acids. The highest concentration of each component is equivalent to 40% of what is typically present in ASM. FICI values are reported as mean ± SEM (n = 3). L) MIC assay for daptomycin against USA300 performed in MHB or ASM with 1/4x, 1x, or 4x the typical mucin concentration, shown as mean ± SEM, n = 6 from three independent experiments.

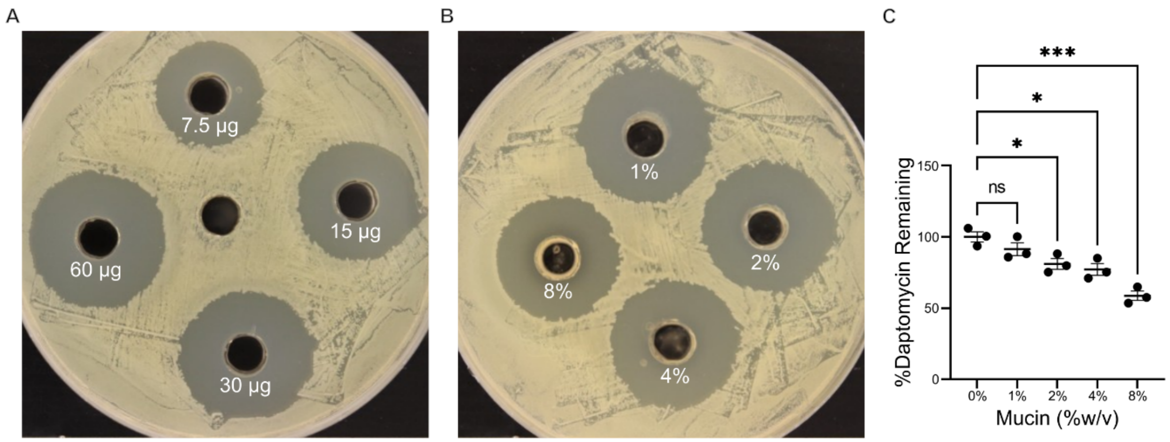


**Supplementary Figure 10. Mucin binds to daptomycin.** Well diffusion antibiotic bioassays performed using USA300 on MHB agar with A) a dilution series of 7.5 µg, 15 µg, 30 µg, and 60 µg daptomycin or B) a fixed dose of 60 µg of daptomycin mixed with a gradient of 1%, 2%, 4%, or 8% w/v mucin. 2% mucin is equivalent to the concentration of mucin present in ASM. C) The concentration of daptomycin remaining following exposure to mucin, calculated using simple linear regression standard curve constructed from A, shown as mean ± SEM, n = 3 from three independent experiments. Significant differences were identified using two-way ANOVA and the Tukey post hoc test. p<0.0001 (****), p<0.001(***), p<0.01(**), and p<0.05(*).

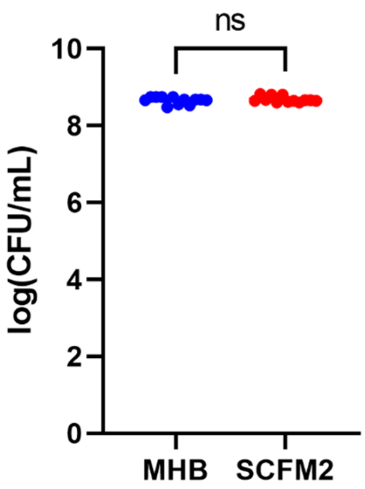


**Supplementary Figure 11. Cell isolation protocol successfully harvests cells from SCFM-2.** CFU counts from USA300 samples harvested from MHB (blue) and SCFM-2 (red). Samples were normalized to an $OD_{600}$ of 1 and drop-plated, incubated, and CFUs were counted. n = 12 from three independent replicates, shown as mean ± SEM. Significant differences were identified using Welch's t-test. p<0.0001 (****), p<0.001(***), p<0.01(**), and p<0.05(*).

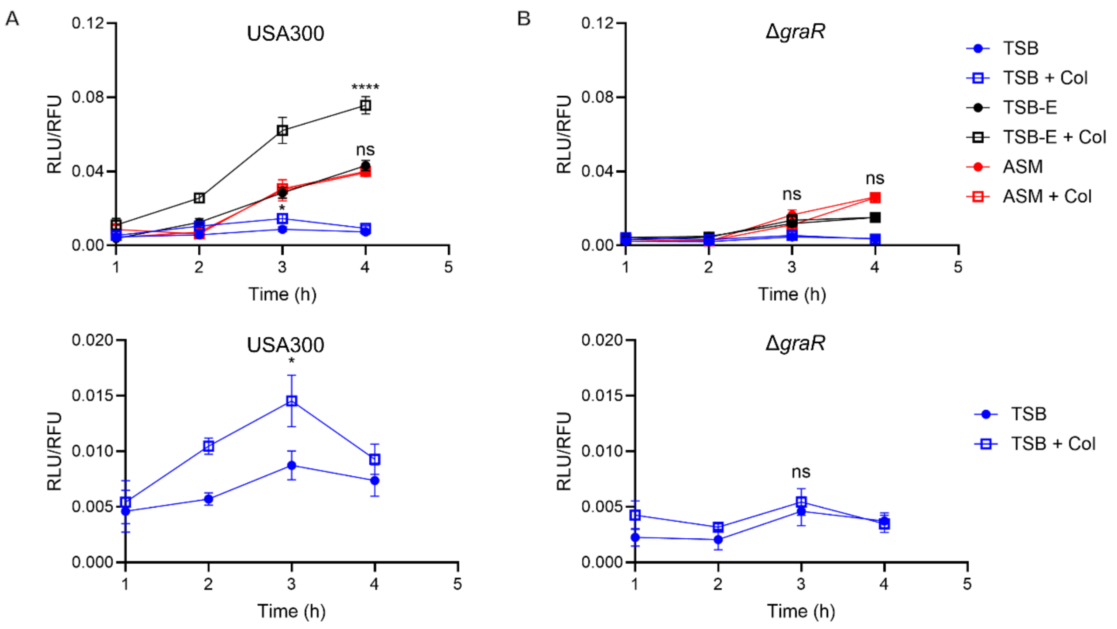


**Supplementary Figure 12. Egg yolk emulsion increases luciferase expression from pGYLux without impacting the induction of the *mprF* promoter.** Monitoring luciferase production under the control of the promoter for *mprF* in A) USA300 and B) Δ*graR* in TSB (blue), TSB with egg yolk emulsion (TSB-E, black), and ASM (red). n=6 from two independent experiments. Statistical analysis was performed comparing induced to uninduced conditions in each medium using two-way ANOVA and the Tukey post-hoc test. $p<0.0001$(****), $p<0.001$(***), $p<0.01$(**), and $p<0.05$(*).

**Supplementary Table 1.** Strains used in this study

| Bacterial Strain | Description | Reference/Source |
|---|---|---|
| *S. aureus* USA300 | Wildtype strain of *S. aureus*; hypervirulent community-associated MRSA; also known as JE2; parent of NTML | Lab stock |
| *S. aureus* Newman | Human clinical MSSA isolate | Lab stock |
| CMRSA1 | HA MRSA USA600 lineage strain, ST45, CC45, SCC*mec* type II | Lab stock[1] |
| CMRSA2 | HA MRSA USA100/800/New York lineage strain, ST5, CC5, SCC*mec* type II | Lab stock[1] |
| CMRSA3 | HA MRSA strain, ST241, CC8, SCC*mec* type III | Lab stock[1] |
| CMRSA4 | HA MRSA USA200/EMRSA16 lineage strain, ST36, CC30, SCC*mec* type II | Lab stock[1] |
| CMRSA5 | HA MRSA USAA500 lineage strain, ST8, CC8, SCC*mec* type IV | Lab stock[1] |
| CMRSA6 | HA MRSA strain, ST239, CC8, SCCmec type III | Lab stock[1] |
| CMRSA7 | CA MRSA USA400/MW2 lineage strain, ST1, CC1, SCC*mec* type IV | Lab stock[1] |
| CMRSA8 | HA MRSA EMRSA15 lineage strain, ST22, CC22, SCC*mec* type IV | Lab stock[1] |
| CMRSA9 | HA MRSA strain, ST8, CC8, SCC*mec* type II | Lab stock[1] |
| CMRSA10 | CA MRSA USA300 lineage strain, ST8, CC8, SCC*mec* type IV | Lab stock[1] |
| CF MRSA Clinical Isolate | MRSA isolate from a CF patient, a 3-year-old boy from Regina, Saskatchewan, Canada | Lab stock |
| CF MSSA Clinical Isolate | MSSA isolate from a CF patient from Regina, Saskatchewan, Canada | Lab stock |
| *S. aureus* NTML | Nebraska Transposon Mutant Library Screening Array; 1920 sequence-defined | [2] |

| | | |
|---|---|---|
| | non-redundant transposon (Tn) mutants of *S. aureus* subsp. *aureus* USA300 JE2 arrayed in five 384-well microtiter plates | |
| *S. aureus* CMRSA10Δ*tarO* | CMRSA10 strain with a *tarO* deletion | 3 |
| USA300-pGYlux | USA300 strain that contains pGYlux, a promoterless bioluminescent reporter plasmid encoding *luxABCDE* | 3 |
| USA300-pGYlux::*mprF* | USA300 wildtype strain that contains pGYlux with *mprF* promoter upstream of *luxABCDE* | 3 |
| USA300-pGYlux::*ilvD* | USA300 wildtype strain that contains pGYlux with *ilvD* promoter upstream of *luxABCDE* | 3 |
| USA300Δ*graR*-pGYlux::*mprF* | USA300 strain with a markerless *graR* deletion that contains pGYlux with *mprF* promoter upstream of *luxABCDE* | 3 |

**Supplementary table 2.** Chemical composition of media used in the study.

| **Component** | **MHB** | **ASM** | **SCFM-2** | *unit* |
|---|---|---|---|---|
| $MgCl_2$•6H2O | 0.12 | - | 0.606 | mM |
| $CaCl_2$ | 0.2 | - | 1.754 | mM |
| Starch | 1.5 | - | - | g/L |
| Beef Extract | 2 | - | - | g/L |
| Casein | 17.5 | - | - | g/L |
| Egg yolk emulsion | - | 0.005 | - | %v/v |
| DTPA | - | 0.015 | - | mM |
| Salmon sperm DNA | - | 1.5 | 0.6 | g/L |
| Bacto CASamino acids | - | 5 | - | g/L |
| BSA | - | 10 | - | g/L |
| mucin | - | 20 | 5 | g/L |
| KCl | - | 29.51 | 14.87 | mM |
| NaCl | - | 85.56 | 51.81 | mM |
| N-acetylglucosamine | - | - | 0.0003 | mM |
| $Na_2HPO_4$•12$H_2O$ | - | - | 0.00125 | mM |
| DOPC | - | - | 0.001272 | mM |
| $NaH_2PO_4$ | - | - | 0.0013 | mM |
| $FeSO_4$•7$H_2O$ | - | - | 0.0036 | mM |
| tryptophan | - | - | 0.013 | mM |
| cysteine•HCl | - | - | 0.16 | mM |
| arginine•HCl | - | - | 0.206 | mM |
| $K_2SO_4$ | - | - | 0.271 | mM |
| $KNO_3$ | - | - | 0.318 | mM |
| histidine•HCl•$H_2O$ | - | - | 0.52 | mM |
| phenylalanine | - | - | 0.53 | mM |
| methionine | - | - | 0.634 | mM |
| ornithine•HCl | - | - | 0.67 | mM |
| tyrosine•2Na | - | - | 0.802 | mM |
| aspartic acid•$H_2O$ | - | - | 0.828 | mM |
| threonine | - | - | 1.072 | mM |
| valine | - | - | 1.118 | mM |
| isoleucine | - | - | 1.12 | mM |
| glycine•HCl | - | - | 1.204 | mM |
| serine | - | - | 1.446 | mM |
| glutamic acid•HCl | - | - | 1.55 | mM |
| leucine | - | - | 1.61 | mM |
| proline | - | - | 1.662 | mM |
| alanine | - | - | 1.78 | mM |
| lysine•HCl | - | - | 2.128 | mM |
| $NH_4Cl$ | - | - | 2.28 | mM |
| Dextrose | - | - | 3 | mM |

| | | | | |
|---|---|---|---|---|
| L-Lactic Acid | - | - | 9.3 | mM |
| MOPS | - | - | 10 | mM |

### Supplementary Methods

**Resazurin reduction assay.** The resazurin cell viability assays were performed using protocols adapted from a previously established method[4]. A resazurin dye solution (Sigma-Aldrich) was added to bacterial cultures at endpoints at a final concentration of 0.014 mg/mL. The plates were then placed in a shaking incubator at 37°C (600 RPM) for 3 hours, ensuring the contents of the plate were not exposed to light. Bacterial growth was measured as relative fluorescence units (RFU) at 560 nm excitation/590 nm emission.

**Isolation of bacteria from liquid cultures.** Cultures were initially pelleted by centrifugation at 200×g for 5 minutes, then resuspended in an equal volume of phosphate-buffered saline (PBS). Subsequently, suspensions were repeatedly centrifuged at 200xg until mucin was sufficiently removed. Cells were then pelleted and resuspended in 1 mL PBS for use in assays. To validate this method, samples were diluted to 1 OD and drop-plated onto LB agar. No difference was observed between isolates from MHB and SCFM-2 (Supplementary Fig. 11).

**Penicillin binding protein (PBP) quantifications.** 30 µg of membrane fractions isolated from USA300 and *pbp4*::Tn grown in either MHB or SCFM-2 were mixed with Bocillin FL at a final concentration of 20 µM and incubated at 37ºC for 10 minutes. Laemmli buffer was then added to the samples, which were then boiled for 5 minutes and separated on a 10% SDS gel for 30 minutes at 80V (or until the samples entered the resolving gel), followed by 135 minutes at 120V. Gels were then visualized using a Typhoon FLA 7000 gel scanner (GE Healthcare). For Western blots, 10 µg of membrane fractions isolated from USA300 and *pbp2a*::Tn grown in either MHB or SCFM-2 were separated on a 10%

SDS gel, as with the PBP quantification protocol. Proteins were transferred to nitrocellulose film using a Mini Trans-Blot® Cell (Bio-Rad) at 100V for 2 hours. The membrane was then blocked overnight at 4ºC with rocking in Blocker™ FL (Thermo Scientific). The primary antibody, mouse anti-*S. aureus*-PBP2a (Thermo Fisher Scientific, diluted 1:1000 in Blocker™ FL) was incubated with the membrane for 2 hours at room temperature with rocking. Horseradish peroxidase conjugated sheep anti-mouse IgG (Thermo Scientific, diluted 1:1000 in Blocker™ FL) was then incubated with the membrane for 2 hours at room temperature with rocking. Finally, the membrane was developed using 1-Step™ Ultra MB-Blotting Solution (Thermo Scientific) for 10 min. Membranes were imaged on a Gel Doc™ EZ Imager (Bio-Rad). Band quantifications were performed using ImageJ.

**Lysostaphin-mediated lysis turbidimetric assay.** Lysostaphin lysis assays were performed as previously described[5]. Briefly, overnight cultures were diluted to a final $OD_{600}$ of 0.05 in 10 mL of fresh MHB or SCFM-2 and grown to an $OD_{600}$ of ~0.6. Cells were then harvested in 1 mL PBS and normalized to an $OD_{600}$ of 0.6. Lysostaphin was subsequently added to a final concentration of 0.5 µg/mL. 200 µL/well of each suspension was then added to a 96-well plate and read every 15 minutes for 3 hours. 200 µL of PBS was used as a blank for background subtraction.

**Cytochrome C binding assay.** Cytochrome C binding assays were performed as previously described, with some modifications[5]. Briefly, cells were grown overnight on solid MHB or solid ASM media (each containing 0.75%w/v agarose added to the respective liquid medium). Cultures were then resuspended in MOPS buffer (20 mM, pH 7.0), normalized to an $OD_{600}$ of 15, and washed twice with MOPS buffer. Cytochrome C

(Sigma-Aldrich) was added at a final concentration of 0.5 mg/mL and incubated for 10 minutes at room temperature. Suspensions were then centrifuged for 5 minutes at 16000 x g, and three 200 µL aliquots of the supernatant were read in a 96-well plate at 530 nm.

**Bioluminescence promoter-reporter assay.** The luciferase expression assays were done as previously described[6], with some modifications. Briefly, overnight cultures of USA300 pGYlux::*mprF*, Δ*graR* pGYlux::*mprF*, USA300 pGYlux::*ilvD*, and USA300 with a pGYlux control vector were grown overnight in tryptic soy broth containing chloramphenicol. Cultures were then diluted to a final $OD_{600}$ of 0.01 in 10 mL of TSB and ASM, with and without colistin, and incubated at 37 ºC with shaking at 220 RPM. After 1 h, the luminescence was read as relative light units (RLU) in a 96-well white polystyrene plate. Cell density of cultures was measured using the resazurin reduction assay. To account for the additional growth during the 3 hours of incubation with resazurin, cultures were diluted 32-fold prior to resazurin addition, and the final RFU values were multiplied by the dilution factor. RLU values were normalized by their corresponding RFU growth values. Higher RLU was detected in ASM due to egg yolk emulsion in the medium (Supplementary fig. 12), and thus statistical analysis comparing conditions with and without colistin were performed independently in each medium.

**(P)ppGpp quantification.** (p)ppGpp was quantified using PyDPA as previously described[7]. Briefly, cultures were grown in MHB and SCFM-2 containing the desired concentrations of antibiotics to an $OD_{600}$ of ~0.6. Cells were then harvested, washed three times with PBS, and normalized to an $OD_{600}$ of 12.5 in 1 mL PBS. Cells were resuspended in 0.25 mL ice cold methanol and vortexed for 1 minute. Cell debris was pelleted via centrifugation at 16000xg for 10 minutes at 4ºC, and the supernatant was mixed 1:1 by volume with HEPES (1 mM, pH 7.0) to achieve a final concentration of 50% methanol and HEPES. PyDPA was added to a final concentration of 40 μM and three 50 μL aliquots were pipetted into a 384W black plate and read in a plate reader at 344 nm excitation/470 nm emission.

**Antibiotic bioassay by agar well diffusion.** Agar well diffusion assays were done following procedures adapted from previously established disc diffusion techniques[8]. Well contents were prepared in microcentrifuge tubes. These included a dilution series of daptomycin (to construct the standard curve) and a fixed dose of daptomycin with or without a gradient of mucin concentration (pH 7). To achieve a semi-confluent lawn of *S. aureus*, a bacterial suspension with $OD_{600}$ of 0.05 was prepared and 100 μL of the suspension was spread evenly across an MHB agar plate and left to dry for 5 min. Five wells were then punched out of the agar for each plate using a sterile P1000 tip and 200 μL of the contents of each microcentrifuge tube was dispensed into the wells. The plates were then placed in an incubator at 37°C for 20 hours. The zone of inhibition (mm) was measured for each well, and the remaining daptomycin concentration of the mucin-containing wells was then interpolated based on the standard curve, calculated using simple linear regression on Graphpad Prism 9.

### References

(1) Christianson, S.; Golding, G. R.; Campbell, J.; Mulvey, M. R. Comparative genomics of Canadian epidemic lineages of methicillin-resistant Staphylococcus aureus. *J Clin Microbiol* **2007**, *45* (6), 1904-1911. DOI: 10.1128/jcm.02500-06 From NLM.

(2) Fey, P. D.; Endres, J. L.; Yajjala, V. K.; Widhelm, T. J.; Boissy, R. J.; Bose, J. L.; Bayles, K. W. A genetic resource for rapid and comprehensive phenotype screening of nonessential *Staphylococcus aureus* genes. *mBio* **2013**, *4* (1), e00537-00512. DOI: 10.1128/mBio.00537-12 From NLM Medline.

(3) El-Halfawy, O. M.; Czarny, T. L.; Flannagan, R. S.; Day, J.; Bozelli, J. C., Jr.; Kuiack, R. C.; Salim, A.; Eckert, P.; Epand, R. M.; McGavin, M. J.; et al. Discovery of an antivirulence compound that reverses beta-lactam resistance in MRSA. *Nat Chem Biol* **2020**, *16* (2), 143-149. DOI: 10.1038/s41589-019-0401-8 From NLM Medline.

(4) Riss, T. L.; Moravec, R. A.; Niles, A. L.; Duellman, S.; Benink, H. A.; Worzella, T. J.; Minor, L. Cell viability assays. *Assay Guidance Manual [Internet]* **2016**.

(5) Douglas, E. J. A.; Palk, N.; Brignoli, T.; Altwiley, D.; Boura, M.; Laabei, M.; Recker, M.; Cheung, G. Y. C.; Liu, R.; Hsieh, R. C.; et al. Extensive re-modelling of the cell wall during the development of *Staphylococcus aureus* bacteraemia. *eLife* **2023**. DOI: 10.7554/elife.87026.2.

(6) Flannagan, R. S.; Kuiack, R. C.; McGavin, M. J.; Heinrichs, D. E. Staphylococcus aureus uses the GraXRS regulatory system to sense and adapt to the acidified phagolysosome in macrophages. *MBio* **2018**, *9* (4), 10.1128/mbio. 01143-01118.

(7) Gao, W.; Chua, K.; Davies, J. K.; Newton, H. J.; Seemann, T.; Harrison, P. F.; Holmes, N. E.; Rhee, H.-W.; Hong, J.-I.; Hartland, E. L. Two novel point mutations in clinical Staphylococcus aureus reduce linezolid susceptibility and switch on the stringent response to promote persistent infection. *PLoS pathogens* **2010**, *6* (6), e1000944.

(8) Andrews, J. BSAC standardized disc susceptibility testing method (version 8). *Journal of antimicrobial chemotherapy* **2009**, *64* (3), 454-489.
